# Subspace communication in the hippocampal-retrosplenial axis

**DOI:** 10.64898/2025.12.31.697203

**Authors:** Joaquín Gonzalez, Mihály Vöröslakos, Deren Aykan, Nina Soto, Noam Nitzan, Rachel Swanson, Zhe Sage Chen, György Buzsáki

**Affiliations:** Department of Psychiatry, New York University Grossman School of Medicine, New York, NY, USA; Neuroscience Institute, New York University Grossman School of Medicine, New York, NY, USA; Department of Neuroscience, New York University Grossman School of Medicine, New York, NY, USA; Department of Biomedical Engineering, New York University Tandon School of Engineering, Brooklyn, NY, USA; Center for Neural Science, New York University, New York, NY, USA

## Abstract

The capacity and flexibility of hippocampal circuits for transforming inputs into downstream outputs is fundamental for navigation and memory, yet the circuit-level mechanisms that allow this operation to adapt across experiences remain unknown. We approach this problem by performing large-scale (up to 1024-channel) recordings across the hippocampal–retrosplenial cortex (RSC) circuit in behaving mice, enabling simultaneous access to spiking activity in dentate gyrus (DG), CA3, CA2, CA1, RSC. Based on a linear dimensionality reduction technique known as partial canonical correlation analysis, we identify low-dimensional communication subspaces^1^ between two regions while accounting for measured third-area influences. These subspaces captured distinct input–output transformations in CA1, linking upstream (DG, CA3, and CA2) hippocampal activity to downstream cortical targets (RSC). Iintrinsic firing properties and anatomical location constrained subspace memberships—members were mapped to deep sublayers of the CA3-CA1-RSC axis during both spatial and non-spatial tasks. These subspaces could recombine overlapping neuronal pools to support distinct interareal interactions across changing experiences and brain states. Reactivation patterns of CA1-CA3 subspaces, but not those of CA1-RSC, during post-experience sleep correlated with replay, reflecting a plasticity-stability balance of the input-output transformation in the hippocampal-retrosplenial axis. Our data suggest a model in which hippocampal-neocortical communication reconfigures predetermined circuit motifs to flexibly encode experiences.

## Introduction

Communication between the hippocampus and neocortex underlies the consolidation of experiences into long-term memories^2–6^, yet the circuit mechanisms governing this exchange remain poorly understood. The hippocampus can be conceived as a side loop of the neocortex, connecting numerous cortical and subcortical brain structures^7,8^. Generally, communication between brain regions is determined by inter-regional connectivity and local neuronal dynamics, together constraining the temporal evolution of neural population activity^9–14^. In most brain regions, including the hippocampus, only a fraction of neurons receive inputs from upstream areas at any given time, and only a fraction send outputs downstream. Whether these groups overlap, and how their activity is coordinated, remains largely unknown.

Effective communication requires local population dynamics to engage the appropriate input-recipient neurons while driving the relevant downstream-projecting neurons. This process can be described mathematically as aligning neural activity between regions within a shared low-dimensional *communication subspace*, through which information flows selectively^1,15–19^. The input–output transformation is then a geometric mapping that converts the input subspace into an output subspace. Characterizing these communication subspaces reveals the neuronal subpopulations engaged in task-relevant interactions, offering insights into the mechanisms underlying circuit operations^16^. While single brain regions have been extensively studied, behavior emerges from coordinated activity across communicating subspaces of many interconnected areas^1,20^. Identifying these communication subspaces requires large-scale recording methods that can monitor sufficiently large populations of neurons in the interconnected regions. Recording from two or more regions simultaneously has become a standard only recently^21–23^. While two-region experiments can provide valuable insights into cross-circuit interactions, they fall short in revealing how the input subspace (linked to an upstream structure) is transformed into the output subspace that supports downstream communication.

To address this gap, we recorded simultaneously from the dentate gyrus (DG), CA3, CA2, CA1, and retrosplenial cortex (RSC) using high-density silicon probes in behaving mice. We applied partial canonical correlation analysis (pCCA) to identify communication subspaces that mediate these inter-regional interactions while controlling for activity in other areas. This approach enabled us to track how CA1 integrated upstream hippocampal inputs and routed selected outputs to the neocortex during wakefulness and sleep. We find that CA1 flexibly reconfigures a shared pool of neurons to integrate distinct inputs from the dentate gyrus, CA3, and CA2, and then selectively engage its RSC targets, providing a context-dependent interface between hippocampus and cortex. These subspaces persist across different mazes but express spatial context selectivity, while neuronal membership in these subspaces is constrained by intrinsic excitability and laminar position within cell body layers. Finally, we show that CA1 input- and output-related subspaces are differentially reactivated during post-task non-rapid-eye-movement (NREM) sleep, especially during hippocampal sharp-wave ripples^24^.

## Results

### Identifying input and output hippocampal subspaces

We performed two complementary experiments to reveal how neuronal populations across different hippocampal fields interact with each other and with the neocortex: 1) head-fixed awake recordings of spontaneous activity, using a 1024-channel SiNAPS probe covering CA1, CA2, CA3, DG regions (n = 3 sessions from 2 mice; Fig. 1a) and 2) freely behaving recordings during novel maze learning, targeting the mouse hippocampus CA1, CA3 and layer 5/6 RSC (Extended Data Figs. 1 and 2), using Neuropixels 2.0 probes (n = 12 sessions from 4 mice; Fig. 1b). DG, CA3, CA2, and CA1 cells were classified as pyramidal cells, narrow and wide interneurons based on the spike waveform features. RSC cells were classified as narrow interneurons and pyramidal cells. Only the putative pyramidal neurons were included for the initial analysis; see Methods for neuron type classification.

**Figure 1.**
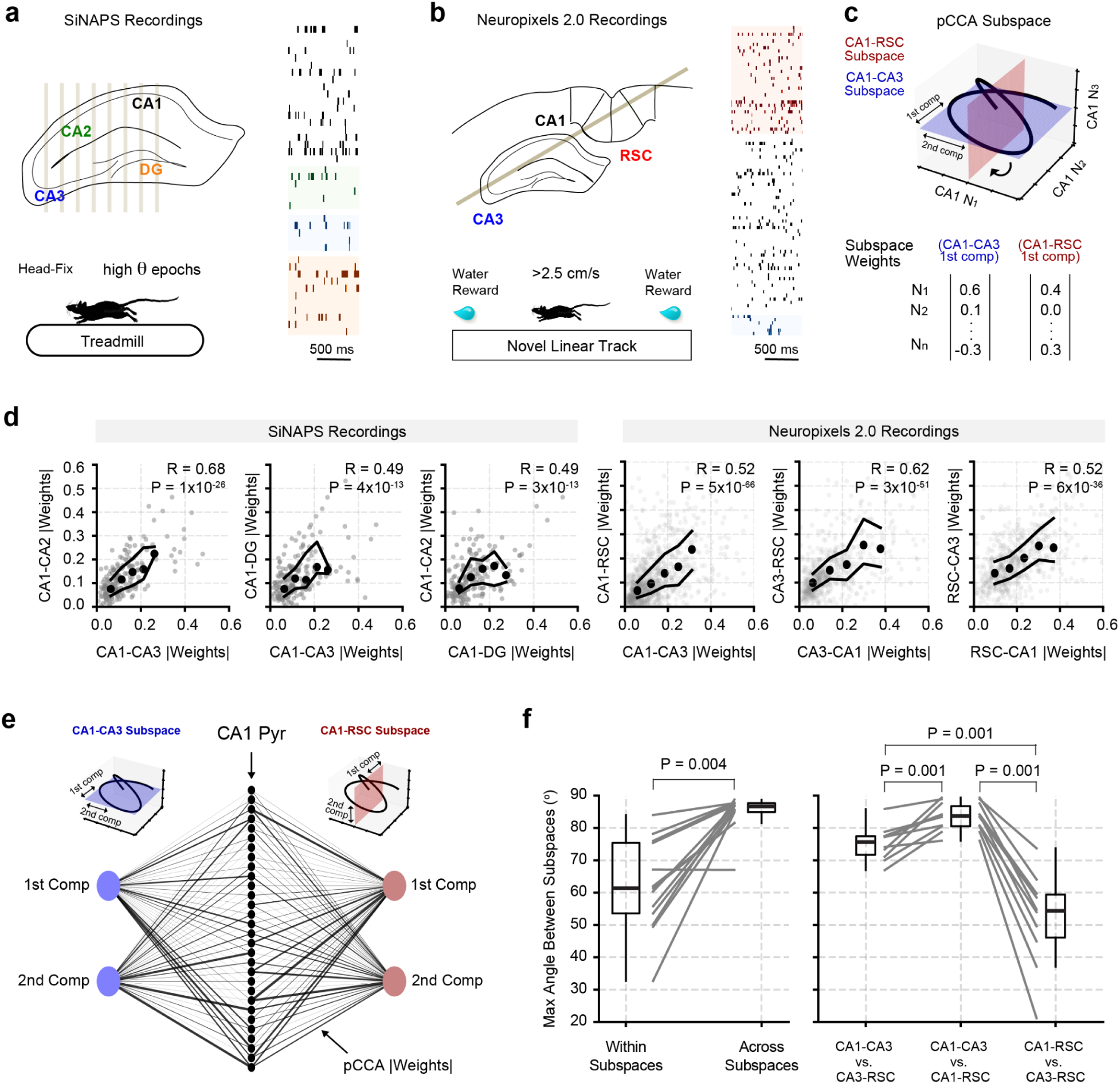
Subspace relationships reveal structured routing and rotation between hippocampal and cortical regions. **(a)** Schematic showing SiNAPS probe targeting hippocampal subregions CA1, CA2, CA3, and DG. Right: example simultaneous recording from all regions. Only pyramidal cells are plotted. **(b)** Schematic showing Neuropixel 2.0 probe targeting CA3, CA1, and layer 5/6 RSC. Right: example simultaneous recording from all regions. Only pyramidal cells are plotted. **(c)** Illustration of pCCA identifying shared subspaces between regions. The solution yields linear combinations of neural activity (weights) in each region that maximize correlation. **(d)** Scatter plots showing for each cell its maximum absolute pCCA weight (across all the significant subspaces it participates). Dark dots represent the median at selected bins, and two solid curves represent 25% and 75% quantiles. Note the strong Pearson correlation (all R statistics were highly significant, P<10^-20^), indicating that the same neurons tend to participate in multiple hippocampal subspaces. **(e)** Example absolute subspace weights from the input (CA1-CA3) and output (CA1-RSC) subspaces in CA1. A total of 30 CA1 pyramidal cells and 2 significant components from each subspace (Dim1, Dim2) are shown. The thickness of the edges shows the weight magnitude, **(f)** Maximum principal angle between distinct subspaces (e.g., CA3–CA1 vs. CA1–RSC). For control in the left panel, the maximum principal angle was computed within each subspace by dividing the trials in half and calculating pCCA independently. *Left panel*: each dot represents the result from one session (n = 13 sessions from 6 mice, combining SiNAPS and NP experiments). For within subspace angles, the results from all within subspaces were averaged to yield a single angle. The same was done for the across subspace group. *Right panel*: populations across areas were matched for the number of neurons. Average rotations across 100 unit-matching samples per session are shown. N = 10 sessions in 4 mice for CA3 vs. CA1 or vs. RSC comparisons. N = 12 sessions in 4 mice for CA1 vs. RSC comparisons. P values were computed from the Wilcoxon signed-rank test.

We applied pCCA to identify linear combinations of neurons that maximized interregional correlations at fast time scales (∼10 ms), while statistically removing the influence of other recorded populations (e.g., eliminating the potential bias of CA3 activity on CA1–RSC correlations, Fig. 1c). Fast timescales were defined by the spike bin width (0.8 ms) and Gaussian smoothing (200 ms kernel, 2.5 ms STD); results were robust across a wide range of bin sizes (Extended Data Fig. 3a,b). For brevity, we refer to these pCCA-derived linear combinations as *communication subspaces*, or simply *subspaces*. To validate the method, we used a three-area Poisson spiking network simulation. This confirmed that pCCA can (1) recover the specific subsets of cells involved in cross-regional interactions, even when subpopulations differ in firing rates (Extended Data Fig. 4a,d,e), and (2) rule out apparent pairwise interactions that are in fact driven by a third recorded area (Extended Data Fig. 4b,c).

We next examined how individual neurons contribute to multiple distinct communication subspaces defined by their upstream and downstream partners (for example, CA1–CA3 and CA1–RSC; Fig. 1c). This analysis was restricted to statistically significant subspace components—pairs of linear combinations across areas—identified using surrogate controls (Extended Data Fig. 3c-f). Neurons with large weight magnitudes in one subspace tended to have large weights in others as well, yielding positive correlations in subspace contributions across all tested pairs (Fig. 1d,e). This widespread overlap in neuronal membership raises a key question: if many neurons participate in multiple communication streams, how does the hippocampal circuit selectively route information between specific inputs and outputs?

We reasoned that selective communication could arise if different combinations of the same neurons are recruited for distinct sender–receiver interactions (Fig. 1e). Formally, this corresponds to the input and output subspaces being rotated relative to each other. We quantified this rotation by measuring the principal angles between pairs of subspaces (e.g., CA1–CA3 vs. CA1–RSC). Principal angles generalize the concept of the angle between two lines, yielding n values that describe the relative orientation of two n-dimensional subspaces. For instance, if the maximum principal angle equals 90°, then the two subspaces are orthogonal along at least one dimension. Given that subspace weights are correlated at the single-neuron level, orthogonal subspaces would indicate that the same neurons are recombined into distinct patterns of coordinated activity.

As a control, we measured the maximum principal angle within the same subspace when computed from separate portions of the recording (e.g., CA1–CA3 in the first half vs. CA1–CA3 in the second half). These angles were consistently smaller than those measured between CA1 subspaces linked to different partners (DG, CA3, CA2, RSC), which approached 90° (Fig. 1f, left). Given that CA1–CA3 and CA1–RSC subspaces exhibited consistent feedforward directionality during maze behavior (CA3 → CA1 → RSC; Extended Data Fig. 5), this stronger rotation between input and output subspaces supports the idea that CA1 can selectively gate communication channels by reconfiguring the coordination patterns of the same neurons.

For cross-region comparisons, we analyzed only Neuropixels 2.0 recordings, which provided high-yield, simultaneous coverage of CA3, CA1, and RSC. To control for sampling differences, we matched the number of neurons from each region within a session. CA1 exhibited significantly larger subspace rotations than either CA3 or RSC (Fig. 1f, right), and CA3 rotations were themselves larger than those in RSC. This gradient suggests that hippocampal circuits, particularly CA1, have a greater capacity than RSC to reconfigure overlapping neuron pools into distinct co-activity patterns, supporting flexible routing of information across the network.

### Subspace rotations support context-dependent communication

We hypothesized that neuronal recombination may account for context-dependent communication, even when circuits are constrained to use the same neurons under different conditions. To test this, we conducted a two-maze generalization experiment while simultaneously recording from CA3, CA1, and RSC populations, tracking the same three-area populations across both mazes. Mice explored two mazes in different spatial contexts (Extended Data Fig. 6a), each offering water rewards at both ends of the track and with approximately the same number of trials, separated by less than five minutes (Fig. 2a, top panel).

**Figure 2.**
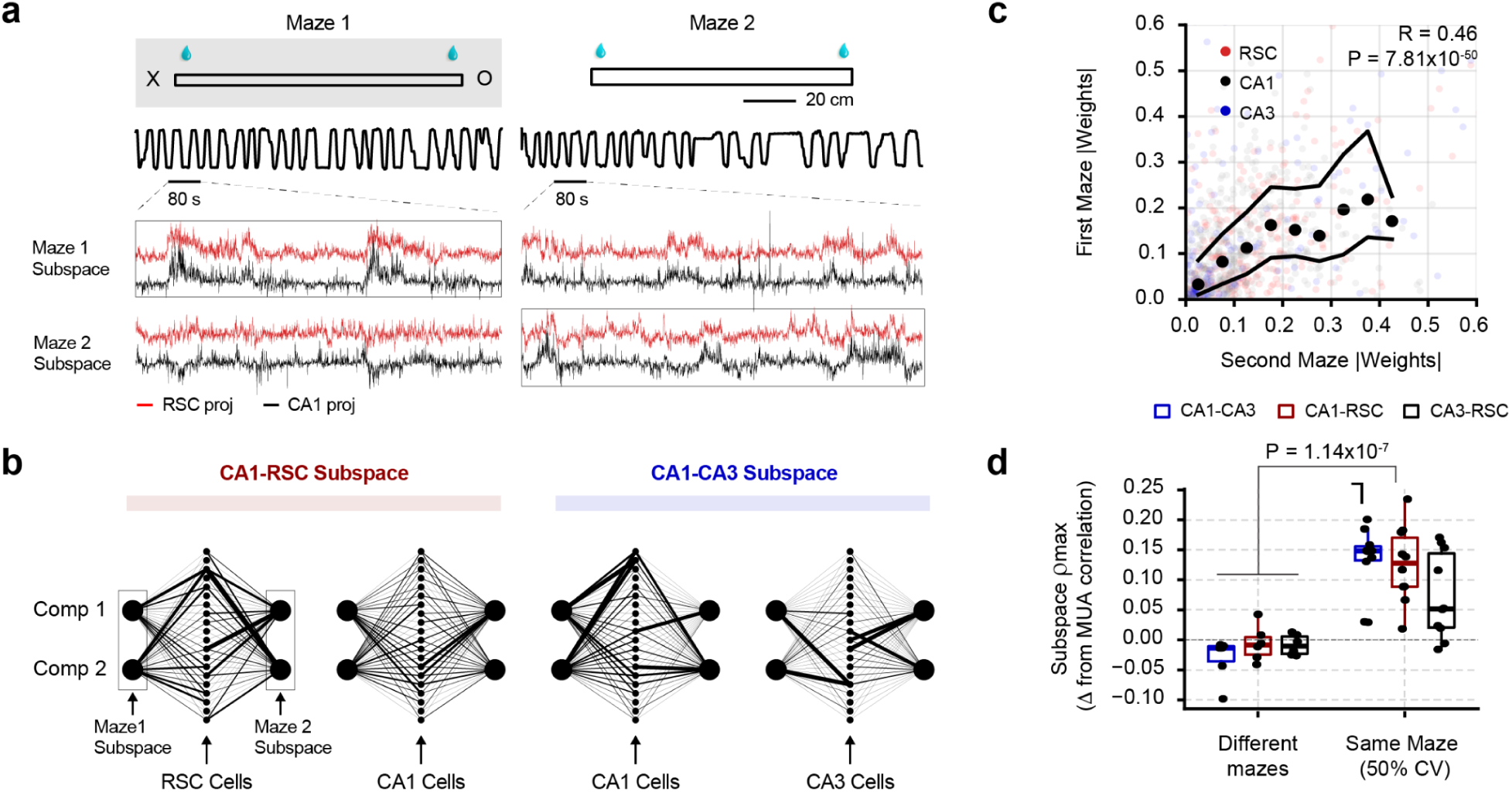
Cross-regional subspaces adapt to distinct environments. **(a)** Schematic of the behavioral protocol. Mice performed in Maze 1 for 30 minutes, followed by a <5-minute intermission, and then explored Maze 2 for an additional 30 minutes. Maze 1 was located within an enclosure in the recording room containing large X and O letters as distal cues. Linearized position trajectory over time is plotted below maze drawings. Traces below show the subspace projected for each area. Subspaces were computed independently on maze 1 (left panels) or maze 2 (right panels). Red and black traces show the subspace activity as a function of time projected on RSC (red) and CA1 (black). **(b)** Example absolute subspace weights from the two mazes in each area. A subset of 50 neurons and 2 significant components from each subspace (Dim1, Dim2) are illustrated. The thickness of the edges shows the magnitude of the weight. **(c)** Single neuron maximum subspace weight magnitudes in CA1, CA3, and RSC were correlated between the two mazes (Pearson correlation R=0.46). **(d)** Subspace correlation ρ_max_ generalization from Maze 1 to Maze 2 or between the first and second half of the trials (Same Maze). All subspaces were computed under one condition and projected to the other. The difference to the MUA correlation is plotted. For the different mazes condition, each dot corresponds to the projection of the subspace computed in one maze to the other (n = 6: two subspaces per session (subspace maze 1 -> maze 2, subspace maze 2 -> maze 1), 3 sessions from 3 mice). For the same maze condition n = 12 sessions from 4 mice, subspace correlation between first and second half (from the subspace computed on the remaining half) were averaged for each session. CV: Cross validation. P values were computed from the Wilcoxon ranksum test.

Despite the expected remapping between mazes^25,26^ (Extended Data Fig. 6b,c), firing rates and the total amount of information transferred through each maze communication subspace remained constant across both environments (Extended Data Fig. 6d,e). By conducting pCCA separately for each maze, we found that neurons across the three regions maintained highly correlated subspace weight magnitudes across contexts (Fig. 2b,c). However, subspace remapping emerged since the combinations employed in each subspace changed from maze 1 to maze 2 (Fig. 2b). Temporal traces in Fig. 2a show the result of computing pCCA on one maze and projecting the weights onto the spiking activity from either the same maze or the different maze. When we projected the subspace computed in one maze onto the other, correlations significantly decreased compared to within-maze correlations and were indistinguishable from those observed in multiunit activity (Fig. 2d). For within-maze comparison, we used 50% cross-validation and computed pCCA on half of the trials and projected it onto the remaining trials, to match the conditions of the two-maze experiment. Importantly, subspace weights were not determined by place field locations (Extended Data Fig. 6f-h, indicating that subspace remapping cannot be solely attributed to place remapping. These results demonstrate that while the same subpopulations of hippocampal neurons are active across different mazes, the combination of coactive cells (i.e., communication subspaces) varies between contexts.

### Subspace membership correlates with firing patterns and anatomical location

The finding that the same neurons remain subspace members across contexts suggests distinct features from their non-member partners. To compare the same number of subspace ‘member’ and ‘non-member’ neurons for fair statistics, we empirically defined “members” as neurons with absolute pCCA weights above the 75th percentile in the top CA1–CA3 and CA1–RSC subspaces, and “non-members” as those below the 25th percentile (Fig. 3a). We focused on the first subspace dimension since it accounted for the most significant interregional correlation (see Methods) and generally explained more of the local variance compared to the other dimensions (Extended Data Fig. 7a). Similar results were obtained using different thresholds or when including all significant subspaces (Extended Data Fig. 7b,c). We analyzed intrinsic properties of individual cells during pre-maze NREM sleep, thereby eliminating the potential confounding effects of learning during maze sessions.

**Figure 3.**
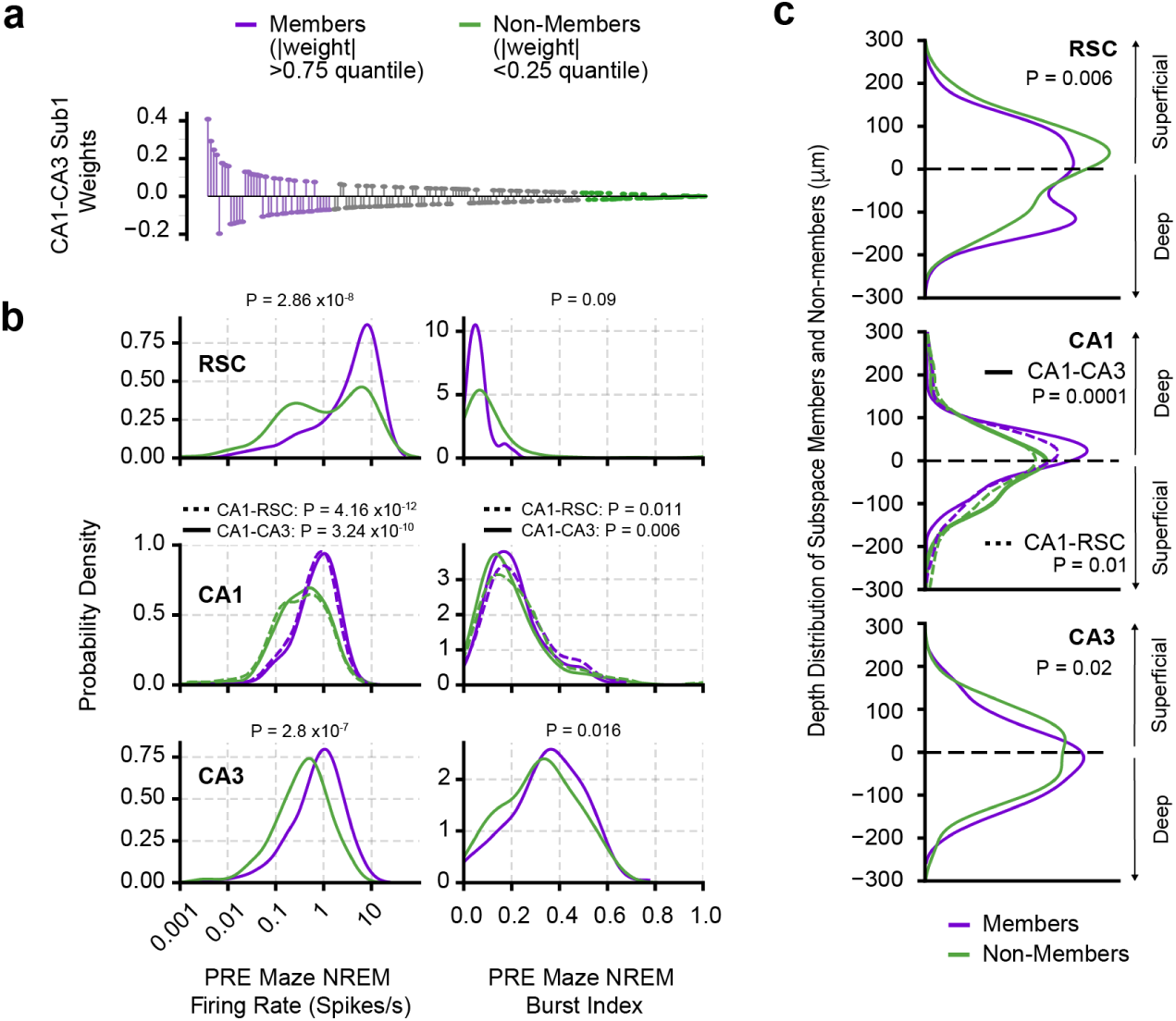
Physiological features of individual neurons predict subspace memberships. **(a)** CA1–CA3 subspace weights (best subspace, the pCCA component accounting for the strongest interregional correlation) for individual neurons. Subspace “members” (violet) are defined as neurons with absolute weights above the 75th percentile of the distribution (|weight| > 0.75 quantile), while non-member neurons (orange) are defined as having absolute weights below the 25th percentile (|weight| < 0.25 quantile). The dominant subspace (accounting for the largest correlations) was employed for this analysis. Similar results are obtained when using all significant subspaces (Extended Data Fig. 7). **(b)** Spiking characteristics of subspace members during pre-maze NREM. Left: Firing rate distributions (log scale) across RSC, CA1, and CA3. Right: Burst index distributions for the same areas. **(c)** Anatomical distribution of subspace members. Depth distribution of subspace members (blue) and non-members (orange) relative to each region’s midplane (defined by the average vertical position of all recorded pyramidal cells in that shank). P values were computed from the Wilcoxon ranksum test.

Member neurons fired at significantly higher rates than non-members across all areas (Fig. 3b). In the hippocampus, but not in RSC, subspace members exhibited more bursts, defined as events with interspike intervals <6 ms (Fig. 3b; Extended Data Fig. 7). These differences reflect preserved propensities of individual neurons, irrespective of waking experience. First, all firing rates were z-scored before computing subspaces, ensuring that all cells had the same mean and standard deviation. Second, the firing rates were measured during pre-maze NREM sleep. Third, subsampling spikes related to different quantiles yielded similar subspace weight magnitudes (Extended Data Fig. 8), indicating that membership is not a consequence of an arbitrary threshold. Finally, our model shows that pCCA identifies truly interacting cells despite them having a lower overall firing rate compared to non-interacting cells (Extended Data Fig. 4e), demonstrating robustness to firing rate biases.

Because firing rates and burstiness also depend on the circuit embeddedness of the neurons^27^, we examined the relationship between subspace membership and sublayer position of the neuronal somata of pyramidal cells. Membership probability was higher for neurons located in deeper sublayers in all three regions (Fig. 3c, Extended Data Fig. 7c^28^). These findings support the hypothesis that subspace membership is not randomly assigned but constrained by preexisting physiological and anatomical properties, suggesting a role for both intrinsic and circuit-level features in shaping interregional communication.

### CA1 subspace supports navigation and other operations

Because computation in hippocampal circuits is most often tested in spatial navigation^8^, we examined the spatial features of pyramidal neurons during a novel linear maze navigation task. Subspace members exhibited temporally structured and coordinated firing patterns aligned with the animal’s movement trajectory. In contrast, non-members displayed less temporally organized activity (Fig. 4a). UMAP^29^ dimensionality reduction of population activity revealed that subspace members preserved the spatial and temporal structure present in the full CA1 manifold, while non-members did not display spatially or temporally organized structure (Fig. 4b,c). Similar results were obtained when defining members across all significant subspaces (Extended Data Fig. 9a,b).

**Figure 4.**
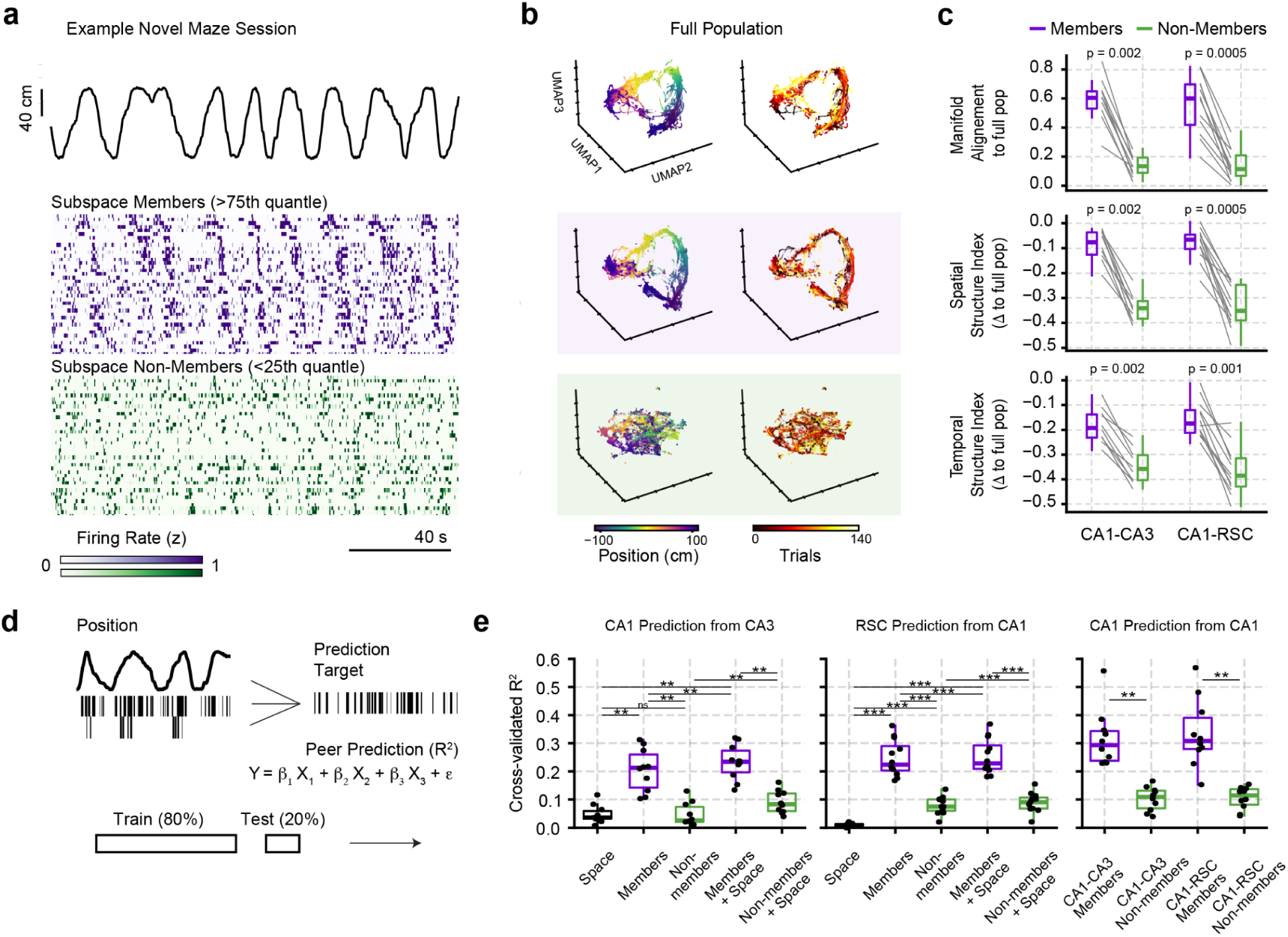
Subspace members preserve maze-related manifold geometry better than non-members. **(a)** Top: Linearized position trajectory over time during a novel maze session. Middle: Z-scored firing rate heatmap for CA1 neurons classified as subspace members (purple) of the CA1–RSC subspace. Units were sorted based on the place field location. Bottom: Firing rates for an equal-sized subset of CA1 non-members (green). The dominant subspace was employed for this analysis. Similar results hold for all significant subspaces. 0.75 and 0.25 quantiles were chosen as thresholds for member and non-member definitions, respectively. Subspace members show more structured and coordinated activity over time compared to non-members. **(b)** 3D UMAP embeddings of population activity during the maze session. Color indicates position (left) or trial number (right). Top: Full CA1 population; Middle: CA1 members in CA1–CA3 subspace. Bottom: CA1 members in CA1–CA3 subspace. **(c)** Top: Manifold alignment score between UMAP embedding of members and non-members to the UMAP embedding of the full population (similarity between the UMAP embedding of the subset and the full population, see Methods). Middle: Spatial structure index quantified over the UMAP embeddings for members and non-members. Bottom: Temporal structure index quantified over the UMAP embeddings for members and non-members. P values were computed from the Wilcoxon signed-rank test. **(d)** Peer prediction schematic, a linear regression model was built to predict the activity of a downstream neuron based on the activity of an upstream neuron and/or the position of the animal along the track. 20% of the data was withheld from training and used to generate a cross-validated prediction. **(e)** Peer prediction R^2^-statistic results averaged for each session. **, P<0.01, ***, P<0.001, Wilcoxon signed-rank test.

While firing rates correlate with several other variables of spatial activity (e.g., the size and number of place fields ^30^), rate differences alone cannot explain our findings. First, firing rates were z-scored prior to the UMAP transformation. Second, subsampled spike trains of members still showed better alignment and higher spatiotemporal structure than those of non-members (Extended Data Fig. 9c,d). Finally, the normalized place fields of members and non-members were similar along the track (Extended Data Fig. 10a,b,c), but differed in their stability; subspace members showed significantly more stable place fields within a session compared to non-members (Extended Data Fig. 10d,e).

We reasoned that neurons with similar place fields can still differ in how stably they contribute to population coding if they vary in their temporal coordination with other cells in the network, potentially explaining why “member” neurons drive population representations more strongly than non-members despite comparable spatial tuning. To test this, we used *peer prediction*^31^ to assess how well the spike timing of downstream subspace members (e.g., CA1) could be predicted from upstream subspace or non-subspace populations (e.g., CA3; Fig. 4d). Subspace members of the upstream area predicted spike times of the downstream member neurons better than non-members and importantly better than space alone (Fig. 4e). Within CA1, subspace members also predicted each other’s activity better than non-members predicted one another (Fig. 4e, right).. Together, these results suggest that a small fraction of inter-regionally connected neurons (i.e., subspace members) drive low-dimensional representations of task structure during behavior.

We next asked whether the deep-layer bias of cross-regional subspace members exclusively reflects the spatial nature of the task or, instead, represents a more general motif. Towards that goal, we analyzed a large-scale visual-coding dataset from the Allen Institute (Extended Data Fig. 11). This dataset includes large-scale simultaneous neuronal recordings from mouse CA3, CA1, and visual areas (primary and higher order), enabling us to assess subspace membership during a non-spatial task (Extended Data Fig. 11a). We confirmed that our spatial navigation-based findings were replicated in this independent dataset: CA1 subspace weights were significantly correlated between CA3- and V1-related subspaces (Extended Data Fig. 11b), and that CA1–CA3 and CA1–V1 subspaces exhibited a significant rotation (Extended Data Fig. 11c). We then examined the laminar position of CA1 neurons by identifying the middle of CA1 pyramidal layer by ripple-associated current source density (CSD) profile (Extended Data Fig. 11d). Deep CA1 cells showed significantly larger CA1–CA3 subspace weights compared to superficial cells (Extended Data Fig. 11e). Importantly, this deep cell-weight bias was absent for CA1–V1 and other higher-order visual area subspaces (Extended Data Fig. 11e), implying that deep-layer CA1-RSC communication is more relevant to spatial navigation.

### Sleep replay of learning-induced subspaces

The repeated activation of the same sequence of neurons in the maze task is expected to be preserved and reactivated during post-experience sleep, supporting memory consolidation^32–35^. We therefore examined whether subspaces established during spatial exploration were reactivated during subsequent post-maze sleep. To do so, we compared the projected activity of the maze-defined subspace with sleep epochs before and after novel spatial exploration (Fig. 5a). Subspace projections from the best maze subspace (the pCCA component accounting for the strongest interregional correlation) exhibited a significant increase in correlation during post-maze NREM sleep compared to pre-maze NREM sleep (Fig. 5b, similar results were obtained from all significant subspaces, Extended Data Fig. 12a). This correlation enhancement was particularly pronounced during sharp wave-ripple (SPW-R) events in post-experience NREM sleep for the CA1-CA3 subspace, but not for the CA1-RSC. The low correlations observed during pre-maze sleep were not statistically different from zero, except for CA1-RSC during ripples (CA1-CA3 NREM: S(9) = 0.58, P = 0.57; CA1-RSC NREM: S(11) = 1.95, P = 0.07; CA1-CA3 ripples: S(9) = -4.56, P = 0.001; CA1-RSC ripples: S(11) = 2.54, P = 0.02; one-sample t-test against zero).

**Figure 5.**
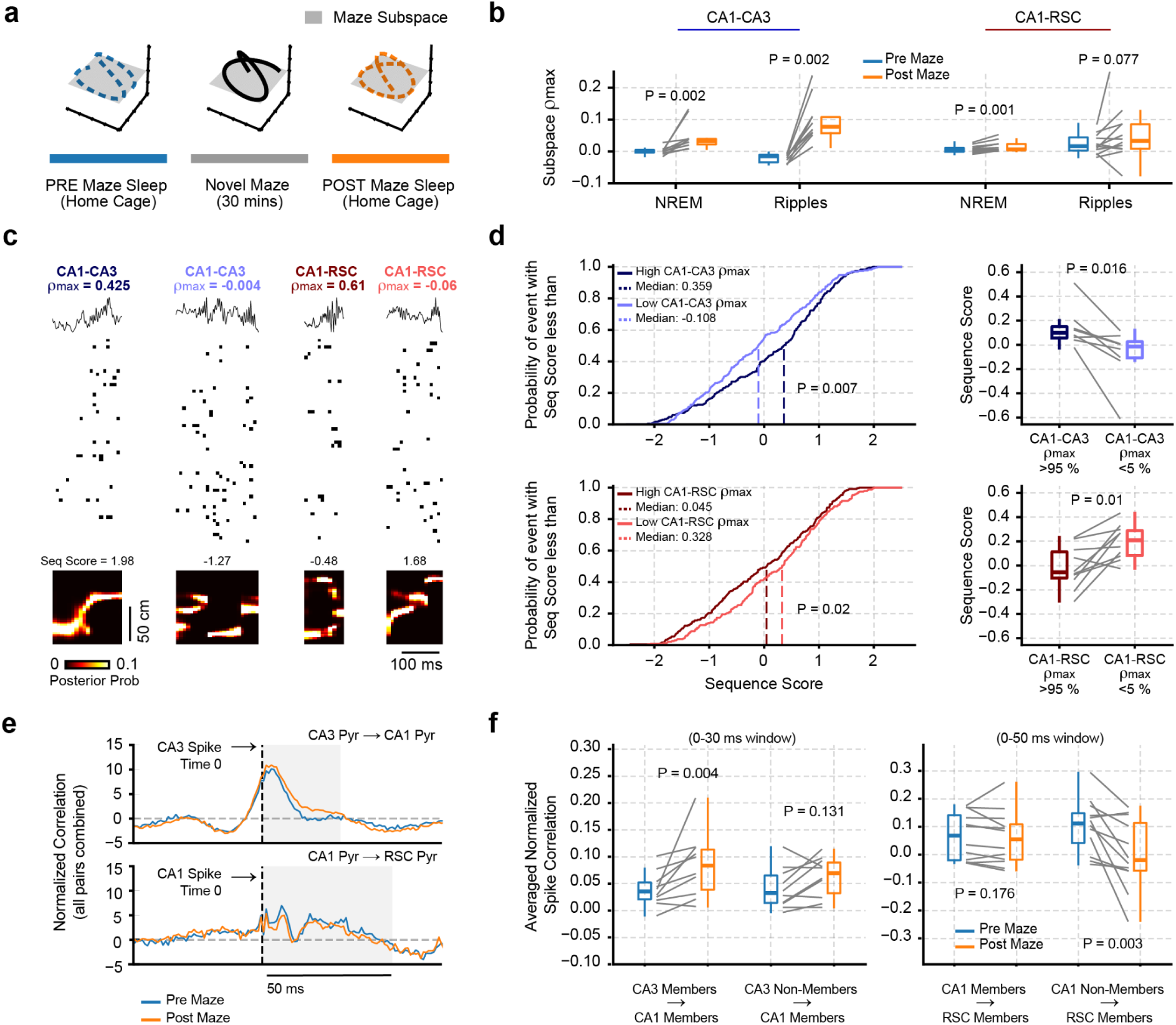
Sleep reactivation of task-defined dominant subspaces predicts replay and spike correlations. **(a)** Schematic showing the projection of maze-defined CA1 subspaces (independently for CA1-CA3 and CA1-RSC) to sleep epochs before (blue) and after (orange) maze exploration. **(b)** Subspace correlations between maze behavior versus NREM and SPW-R events. The best subspace was used for comparisons, but similar results were obtained for all significant subspaces (Extended Data Fig. 12). **(c)** Spike rasters and decoded posterior probabilities for representative SPW-R events^36,37^. values indicate subspace correlations. Note that ρ_max_ for CA1-RSC could be low even when the sequence was regularly ordered (last column). Sequence score was computed for each candidate replay event. P values were computed from the Wilcoxon signed-rank test. **(d)** Left: Cumulative distribution functions of replay scores for SPW-R events with high vs. low subspace correlation values. P values were computed from a Kolmogorov-Smirnov test. Right: Mean sequence scores for each session for high vs. low subspace correlation events. Only the sessions with a median position decoded error <20 cm were included in the analysis. P values were computed from the Wilcoxon signed-rank test. **(e)** Normalized population cross-correlograms. All pairwise cross-correlograms (n = ∼10,000 pairs) were pooled together across all pairs and sessions. Normalization consisted of subtracting the slow cross-correlogram fit and dividing the difference by the number of spikes emitted by the source population (see Methods). **(f)** Spike cross-correlations between pre-maze and post-maze sleep sessions. Each line shows a single session. For each session, the normalized cross-correlograms were integrated in a 0-30 ms (CA3-CA1) or 0-50 ms (CA1-RSC) window (shaded area in panel **e**) and then the results were averaged (n ∼ 1000 correlograms). The same 75th and 25th percentile threshold was employed to define members and non-members. P values were computed from the Wilcoxon signed-rank test

Additionally, we computed subspaces using only pre- and post-maze sleep (Extended Data Fig. 12b). We observed that the subspace weights (for the best maze subspace) were significantly more correlated between pre and post for the CA1-RSC subspace compared to the CA1-CA3 subspace (Extended Data Fig. 12c), suggesting stronger plasticity for intrahippocampal compared to hippocampal-neocortical communication. To understand if new CA1-CA3 neuronal combinations were introduced specifically during maze learning, we quantified the alignment between sleep and maze subspaces. We found that CA1-CA3 subspaces, but not CA1-RSC, were significantly more aligned with the maze subspace during post experience sleep compared to pre-experience sleep (Extended Data Fig. 12d). Therefore, these results suggest that novel experiences drive the formation of new neuronal combinations in the hippocampus.

Because CA1-CA3 subspaces exhibited a higher degree of plasticity, based on the degree of reconfiguration measured, we investigated whether differential subspace reactivations during post-experience NREM sleep bias the replay content. For this analysis, we trained a Bayesian decoder during maze runs (Extended Data Fig. 12e,f) and then decoded ‘virtual trajectories during post-experience sleep epochs (Fig. 5c)^38–41^. Replay events were significantly more likely to occur when the corresponding sleep epochs exhibited high CA1–CA3 subspace correlation (Fig. 5c,d). In contrast, events with high CA1–RSC subspace correlations were associated with reduced replay fidelity compared to low subspace correlation CA1–RSC events (Fig. 5c,d, bottom). This was consistent with the lack of CA1-RSC subspace reactivation during SPW-R and its decreased plasticity compared to CA1-CA3. These findings suggest that distinct inter-regional subspaces differentially contribute to the content and fidelity of reactivated waking sequences.

An important implication of these observations is that the distinct reactivations of CA1–CA3 and CA1–RSC subspaces have different functional consequences. To address this hypothesis, we first computed the normalized population cross-correlograms during the entire pre-experience and post-experience sessions (Methods). While overall CA3-CA1 correlations increased during the post-experience period (0-30 ms), CA1-RSC correlations instead decreased compared to the pre-maze period (0-50 ms, Fig. 5e). To understand how the differential subspace reactivations contributed to these post/pre changes, we computed the average normalized cross-correlogram for each session. We first integrated the normalized correlations for each pairwise cross-correlogram observed in the window at the population level. Neuron pairs were then grouped as either members or non-members in the source region projecting to members in the target region. Spike correlations between CA3 and CA1 pyramidal neurons were selectively enhanced after the maze experience, but only among subspace members (Fig. 5e,f). In contrast, spike correlations in the CA1–RSC subspace did not change among members, but significantly decreased from CA1 non-members to RSC members. These effects were not attributed to changes in firing rate. RSC firing rates remained stable across pre- and post-experience sleep epochs (Extended Data Fig. 12g).

## Discussion

A long-standing puzzle is how very different intrinsic properties of neurons and their anatomical connectivity constraints in the hippocampal regions enable flexible and efficient computation for input-output communication. We described functionally connected subspaces in the DG-CA3-CA2-CA1-RSC network that route context-specific input–output transformations. These low-dimensional subspaces dynamically rotate within selectively shared neuronal populations, allowing different combinations of the same neuronal pools to engage in cross-regional interactions across changing experiences or brain states. Our data suggest a model in which experience builds primarily from a preconfigured reservoir across multiple stages of the hippocampal loop.

Although subspace membership is not a bimodal feature but a continuum, neurons at the two extremes of the distribution can be assigned as either members or non-members. In principle, such subspaces could be flexibly constructed from all neurons in a circuit during particular goals, for example, place cells^8^ and time cells^42^ in spatial and temporal tasks. Alternatively, anatomical constraints can bias how neurons participate in subspace formations. We found that neuron members of the functionally connected subspaces during navigation were anatomically mapped onto the deep layers of the CA3-CA1-RSC axis. This preferential hippocampal subspace connectivity also persisted in a non-spatial, visual experiment.

Subspace weight magnitudes of individual neurons were positively correlated across all tested inter-regional combinations, including DG-CA1, CA3-CA2-CA1, and CA1-RSC. Critically, subspace member neurons across anatomical regions maintained similar subspace weight magnitudes in different contexts. Yet, when the subspace computed in one maze was projected onto another, the correlations were indistinguishable from those observed in multiunit activity, demonstrating that while the same subpopulations of neurons participate across different mazes, the combinations of coactive cells ‘remap’ between contexts. Finally, subspaces also imposed a structure on post-experience plasticity. Task-defined subspaces were reactivated during NREM sleep, particularly during SPW-R events, where they segregated into distinct plasticity regimes. Specifically, hippocampal CA1–CA3 subspaces exhibited enhanced replay and increased spike correlations among their members. In contrast, CA1–RSC subspace activity was associated with reduced replay fidelity and weakened spike transmission from non-member to member cells, suggesting that different plasticity rules might be at work within the hippocampus and across the hippocampus-RSC interface^43,44^.

### Plastic yet stable subspaces at the CA1 interface

Our findings reveal that plastic and stable subspaces can co-exist in the hippocampus, enabling local flexibility embedded within a globally stable communication architecture. Systems consolidation theory and the two-stage model for memory consolidation posit that the hippocampus is an inherently plastic substrate for learning, continuously reconfiguring cells to encode novel spatial and episodic experiences^2–6^. In contrast, retrosplenial and other neocortical networks exhibit more stable, slowly evolving population codes that integrate information over longer time scales^45^. At the interface of these two regimes, CA1 must reconcile the need for rapid context-dependent reconfiguration with the demand for stable transmission to cortical targets. We directly observed this dichotomy: CA1–CA3 subspaces were selectively remodeled by novel maze learning, showing stronger post-maze alignment to the behaviorally defined subspace, enhanced ripple reactivation and trajectory replay. By contrast, CA1–RSC subspaces remained largely stable across pre-experience and post-experience sleep. Moreover, the differential spiking correlation changes between pre and post maze sleep suggest that CA1 does not simply relay their inputs but implements a filtering layer, where intrahippocampal subspaces (CA1–CA3) remain malleable for encoding and consolidation^46,47^, while hippocampal-cortical subspaces (CA1–RSC) preserve a more rigid scaffold optimized for long-term integration^45^.

### Developmentally preconfigured subspaces

Our findings suggest that the distinct propensities of subspace member neurons^48^ and their consistent anatomical bias toward deep layers comprise a preconfigured space, whereas non-members mainly reside in the superficial layers. These selective subspaces are implemented not by recruiting different cell groups across contexts, but by rearranging coactivity patterns within the same population. Such a generalized sparse coding framework is computationally more efficient, as the coactivity patterns can be reshaped to store and retrieve memories with minimal overlap^49–52^. This rotation of subspace geometry across mazes echoes a similar computational principle observed in motor cortex and other systems, where low-dimensional population dynamics evolve along rotated yet constrained trajectories to enable task-specific outputs despite similar inputs^53,54^ ^55^. The functionally defined subspace and non-subspace memberships in the CA1 region mirror the demonstrated differences between ‘deep’ and ‘superficial’ layer neurons^27,56–58^. CA1 superficial and deep layer neurons exhibit prominent radial differences in firing properties, pyramidal-interneuron interactions, afferent and efferent connections, population cooperativity, developmental origin, and behavioral correlates^59–63^ and display superficial-to-deep layer feedforward inhibition^60^. Deep CA1 neurons have larger and more place fields and exhibit increased phase-locking to network oscillations, while superficial cells typically form single but more stable place fields, have higher context specificity, and are specialized in prospective coding^27,57,64^. Similar distinctions have been reported in the radial axis of the CA3 pyramidal layer, including higher firing rates and stronger bursting in the earlier-born, deeper neurons with long apical shafts and sparse mossy fiber innervation^65–72^. In contrast, superficial CA3 pyramidal cells consistently have granule cell inputs (‘thorny’ cells^68^). The distinct nature of the deep and superficial CA3 parallel recurrent networks is further exemplified by their asymmetric excitatory communication and different interneuron targets^73^. Excitatory connectivity is strong from superficial to deep, but sparse from deep to superficial pyramidal cells^73^.

Similar directed superficial-deep asymmetric communication has also been reported in the presubiculum^74^ and the neocortex^75,76^. Thus, the two-layer organization is a general principle that applies to both the hippocampal formation and the neocortex. Both architectures may be based on the same Bauplan, with the neocortex being a more elaborated circuit version of the hippocampus^73^. Our analysis demonstrates that these sublayers maintain their relatively distinct communication streams. These observations shed a novel light on neocortical-hippocampal communication.

### Hippocampal subspaces as a continuation of neocortical “where” and “what” streams

The content of neocortical neuronal activity that flows into the hippocampus via the lateral and medial divisions of the entorhinal cortex (LEC; MEC) is often conceptualized as two parallel streams, traditionally referred to as the ventral (“what”) and dorsal (“where,” “when,” “how”) pathways^77,78^. The dorsal stream is proposed to analyze temporal and spatial sequences of the external world^78,79^and to register and imagine the consequences of actions without the need to overtly execute them^80^. The ventral stream processes items or relations of items based on structure, independent of the moment or position, to extract abstract meaning^79^. The MEC and LEC are anatomical continuations of the dorsal and ventral streams, conveying qualitatively different types of neuronal messages^81–83^. It has been hypothesized that the two streams are mixed and integrated into episodic memory in the hippocampus^42^. This hypothesis aligns with a computational model in which successive parallel and recurrent circuits serially perform pattern separation and pattern completion^84,85^.

Our findings offer an alternative view that the ‘where’ and ‘what’ streams remain relatively segregated at the serially connected hippocampal subfields. MEC and LEC inputs differentially activate dentate granule cells, mossy cells, and CA3 pyramidal cells^83^. In turn, these subcircuits may activate the superficial and deep subspaces we described here. One possible interpretation of this ‘deep-deep-deep’ preferential communication is that the “where,” “when,” “how” functions continue to be selectively processed within the multistage hippocampal system. In turn, the outputs of the deep and superficial CA1 layers route segregated processing to the RSC-prefrontal cortex and the entorhinal cortex, respectively^57^. Although information carried by the “what” and “where” streams can be combined at virtually all levels of the neuraxis, this parallel arrangement allows flexibility for the streams to be kept separate or combined at increasing levels of abstraction. Thus, segregation and integration in the hippocampus may occur not only serially, but also adaptively at each level of the multisynaptic stages. Our analysis of the non-spatial, visually cued responses in the superficial and deep hippocampal sublayers provides support for the dual-stream framework.

Another, although non-exclusive, interpretation of our findings is that the hippocampus utilizes two cooperative yet segregated streams for all its computations. The preserved deep CA3-deep CA1 subspace communication in a non-spatial, visually cued task supports this interpretation. Under this hypothesis, the relatively rigid but more active deep layer scaffold can produce quick ‘good enough’ solutions and generalizations. In contrast, detailed characterization of unique situations, a requisite for episodic memory, may require the engagement of the more flexible, superficial subspaces ^32,57,86,87^, in cooperation with the deep-layer reservoir. Viewed from this perspective, the ‘where’ and ‘what’ streams complement each other, providing a context for the more explicit semantic representations in the superficial chain.

## Methods

### Animal handling

All experimental procedures were conducted in accordance with the National Institutes of Health guidelines and with the approval of the New York University Grossman School of Medicine Institutional Animal Care and Use Committee (IACUC). Mice (adult male C57BL, 26-31 g, n = 4 Neuropixel experiments; n = 2 SiNAPS experiments) were kept in a vivarium on a 12-hour light/dark cycle and were housed 4 per cage before surgery and individually after it. Animals were handled daily and accommodated to the experimenter before the surgery and behavioral recording.

### Surgical procedures

Mice were anesthetized with isoflurane. The body temperature was monitored and kept constant at 36–37 °C with a DC temperature controller (TCAT-LV; Physitemp, Clifton, NJ, USA). Stages of anesthesia were maintained by confirming the lack of a nociceptive reflex. Atropine (0.05 mg kg–1, s.c.) was administered after isoflurane anesthesia induction to reduce saliva production. The skin of the head was shaved, and the surface of the skull was cleaned by hydrogen peroxide (2%). A custom 3D-printed baseplate^88^ (Form2 printer, FormLabs, Sommerville, MA) was attached to the skull using C&B Metabond dental cement (Parkell, Edgewood, NY). A stainless-steel ground screw (90910A310, McMaster-Carr) was placed above the cerebellum, and a craniotomy was performed.

### SiNAPS Probe

For head-fixed experiments^89^, a custom 3D-printed headpost^90^ (Form2 printer, FormLabs, Sommerville, MA) was attached to the skull using C&B Metabond dental cement (Parkell, Edgewood, NY). The location of the craniotomy was marked and a stainless-steel ground screw with a header pin was placed above the cerebellum. Each animal recovered for at least 7 days prior to habituation of the head-fixation. Animals were allowed to walk freely on a treadmill (https://github.com/misiVoroslakos/3D_printed_designs/tree/main/Treadmill_Rinberg) during recording sessions. The day before recording, a craniotomy was performed (2 mm posterior from Bregma and 1.5 mm lateral to midline targeting hippocampus and 2 mm anterior from Bregma and 0.5 mm lateral to midline targeting midline cortices) and the dura was removed. After the surgery, the craniotomy was sealed with Kwik–Sil (World Precision Instruments, Sarasota, FL) until the recording.

On the day of the recording the animal was head-fixed, the craniotomy was cleaned and the headpost was filled with sterile saline. The ground of the probe PCB was connected to the header pin and the probe was inserted into the target depth using a manual micromanipulator (MM-33, Sutter Instruments, Novato, CA). The electrophysiological signal and transistor bias were constantly monitored during insertion. The collected data was digitized at 20 kS s−1 using Smartbox Pro and Activus SiNAPS interface box and visualized using Radiens Allego software (NeuroNexus Technologies, Ann Arbor, MI). At least 30 min were waited after reaching the target depth before starting the data acquisition. A total of 2 h sessions were recorded from each mouse. The 1024 channel SiNAPS probe was employed^89^. Experiments were performed using the commercial system available from Neuronexus implements similar functionalities through the SmartBox Pro and the SiNAPS Interface Box (https://www.neuronexus.com/products/sinaps-interface-box/) which were interconnected through a standard HDMI cable. As for the SiNAPS Research Platform, data from each available electrode-pixel was acquired and digitized at 20 kS s−1, with a 12-bit resolution over a 2.4 dynamic range through the SiNAPS Interface Box which also implemented the same SiNAPS IP core (Corticale, Italy) and outputs electrode-pixel data processed in real-time by the FPGA (i.e., signals filtered over 2 Hz or 300 Hz to 5 kHz). A single HDMI cable was used to transmit in real-time the digitized and filtered data from the SiNAPS BOX Interface to the SmartBox Pro which, in turn, implemented a high-rate data transmission to the PC, where data was stored, analyzed, and visualized by the Allego Software (Neuronexus, USA).

### Neuropixels 2.0

Neuropixels 2.0^22^ was mounted onto a recoverable metal microdrive^88^ and was implanted 2 mm posterior to bregma and 1.1 mm lateral to midline for the contralateral side (Extended Data Fig. 1,2). The probe spanned deep and superficial layers of RSC and area CA1^91^ and CA3 of the hippocampus contralateral to the insertion. The probe was lowered from the surface of the brain at 5 mm DV (Extended Data Fig. 1,2). The craniotomy was sealed with Dura-Gel (Cambridge NeuroTech). The probe and microdrive were then enclosed in a copper mesh built mechanical structure^92^. Each animal recovered for at least 7 days prior to recording. The collected data was digitized at 30 kS/s and a custom PXIe (Peripheral Component Interconnect (PCI) eXtension for Instrumentation; a standardized modular electronic instrumentation platform) data acquisition card was connected to a computer via a PXI chassis (NI 1071, National Instruments, Austin, TX), and OpenEphys or SpikeGLX software was used to write the data to disk^21,93^.

#### Behavioral Training and Experimental Design

Prior to probe implantation, animals were habituated to a linear maze for three consecutive days, with each session lasting 30 minutes. Following habituation, animals were placed on a water restriction schedule and trained to associate water rewards with the linear maze. After recovering from probe implantation surgery, animals were again water-restricted for 24 hours before beginning experimental sessions. Each session began with a pre-sleep period recorded in the animal’s home cage (∼2 hours). Following this baseline, animals performed a novel linear maze exploration. The linear mazes were placed in novel environments (Room A or B), and each maze track differed in shape, color, and construction materials to ensure distinctiveness (Supplementary Table 1). During maze exploration, animals alternated between two water wells located at either end of the track, retrieving rewards for approximately 30 – 60 minutes. After the linear maze run, animals were returned to their home cage, and a post-sleep session was recorded. Mice (n = 4) were tested in 3 novel maze runs (Supplementary Table 1).

Three mice were also recorded in a novel environment using two different mazes. Following the pre-sleep session, each animal was placed in the first maze, which was enclosed by a black room divider featuring white “X” and “O” markings at each end, near the water reward locations. After completing 30 trials, the animal was returned to its home cage. The maze was then replaced with a second one, and the black room divider was removed. The animal subsequently continued running in the new linear maze. After completing the second maze, the animal was again returned to its home cage, and a post-sleep recording session was conducted. While the animal had experienced both mazes previously, the room enclosure and context changed in both scenarios.

Infrared (IR) sensors were used to detect the animal’s run towards a water port. IR sensors and syringe pumps (https://karpova-lab.github.io/syringe-pump/latest/) for water delivery were controlled by a customized Arduino-based circuit. The animal’s position was tracked with an overhead Basler camera (catalog no. acA1300-60 gmNIR, Graftek Imaging) at a frame rate of 25 Hz, and tracking data were aligned to the recording via transistor - transistor logic pulses from the camera, as well as a slow pulsing LED located outside the maze. The animal’s location was detected within a region of interest around the maze, using a custom-trained DeepLabCut neural network^94^. Detections with a likelihood below 0.5 were discarded. The occasionally missing trajectory detections were filled using MATLAB function “fillmissing” with method “pchip” which is a shape-preserving piecewise cubic spline interpolation and were then smoothed using a 7th-order one-dimensional median filter “medfilt1”.

#### Allen Institute dataset

The publicly available dataset was collected from mice in virtual reality experiments at the Allen Institute (Allen Brain Observatory – Neuropixels Visual Coding; https://allensdk.readthedocs.io/en/latest/visual_coding_neuropixels.html). In the “functional connectivity” protocol, the natural stimuli of one 30-s B/W movie (30 frames per second) with semantic information were presented to animals 60 times. The multi-Neuropixel recordings consisted of concurrent electrophysiological data of many brain regions (primary and high-order visual cortex, hippocampal subfields CA1, CA3 and DG).

### Single unit analysis

A concatenated signal file was prepared by merging all recordings from a single animal from a single day). Putative single units were first sorted using Kilosort2.5^94,95^ and then manually curated using Phy (https://phy-contrib.readthedocs.io/).

### Cell Type Classification

In the processing pipeline, cells were classified into two putative cell types: interneurons, and pyramidal cells. Interneurons were selected by two separate criteria. We labeled single units as interneurons if their waveform trough-to-peak latency was <0.425 ms, or if the waveform trough-to-peak latency was >0.425 ms and the rise time of the autocorrelation histogram was >6 ms. The remaining cells were assigned as pyramidal cells. Autocorrelation histograms were fitted with a triple exponential equation to supplement the classical, waveform feature based single unit classification (https://cellexplorer.org/pipeline/cell-type-classification/)^96^. Bursts were defined as groups of spikes with interspike intervals < 9 ms.

#### SPW-R detection and properties

SPW-Rs were detected from manually selected channels located in the center of the CA1 pyramidal layer. Broadband LFP was bandpass-filtered between 130 and 200 Hz using a third-order Chebyshev filter, and the normalized squared signal was calculated. SPW-R peaks were detected by thresholding the normalized squared signal at 5×SDs above the mean, and the surrounding SPW-R begin, and end times were identified as crossings of 2×SDs around this peak. SPW-R duration limits were set to be between 20 and 200 ms. An exclusion criterion was provided by manually designating a ‘noise’ channel (no detectable SPW-Rs in the LFP), and events detected on this channel were interpreted as false positives (e.g., EMG artifacts). The ripple detection quality was visually examined by superimposing the detected timestamps on the raw LFP traces in NeuroScope2 software suite.

#### Sleep state scoring and LFP analysis

Brain state scoring was performed as described in the study by^97^. In short, spectrograms were constructed with a 1-s sliding 10-s window fast Fourier transform of 1,250 Hz data at log-spaced frequencies between 1 Hz and 100 Hz. Three types of signals were used to score states: broadband LFP, narrowband high-frequency LFP and electromyogram (EMG) calculated from the LFP. For broadband LFP signal, principal component analysis (PCA) was applied to the Z-transformed (1–100 Hz) spectrogram. The first principal component in all cases was based on power in the low (32 Hz) frequencies. Dominance was taken to be the ratio of the power at 5–10 Hz and 2–16 Hz from the spectrogram. All states were inspected and curated manually, and corrections were made when discrepancies between automated scoring and user assessment occurred.

### Channel selection for Neuropixels 2.0 recordings

After full recovery from surgery, we recorded a home-cage sleep session with a linear probe configuration to reconstruct the electroanatomy of RSC, CA1 and CA3. We recorded from every 4^th^ channel on each shank simultaneously, resulting in a total of 92 channels per shank. This configuration spanned 5,760 µm along the dorso-ventral axis and 750 µm medio-laterally.

To localize anatomical landmarks, we first detected sharp-wave ripples (SPW-R) in the pyramidal layer of CA1^24^ and constructed an event-triggered average current source density (CSD) map^98^ (Extended Data Fig. 13). The 2-dimensional CSD map confirmed the source-sink-source distribution of SWRs. To further refine the boundaries between hippocampal and cortical regions, we incorporated single unit activity (Extended Data Fig. 13). Neuronal soma locations were estimated based on the peak-to-peak amplitude of spike waveforms^96,99^ (Extended Data Fig. 13) and mapped to the corresponding electrode on each shank. The cell-free zone above the CA1 pyramidal layer corresponded to alveus, the ventricle and the corpus callosum, while the first cellular layer above the corpus callosum was identified as the bottom of RSC. Due to the limited number of simultaneously recordable sites with the Neuropixels probe (384), we limited our RSC recordings to layers 5/6. Thus, when describing “superficial” and “deep” layers of the RSC, were refer to the relative positions of the recorded units, rather than the conventional superficial (layers 1-4) and deep (layers 5-6) of the neocortex. CA3 was identified by the presence of a neuron-dense layer located deeper than the CA1 pyramidal layer, exhibiting synchronous firing during SPW-Rs. Once these anatomical boundaries were defined, we used SpikeGLX (https://billkarsh.github.io/SpikeGLX/) to optimize channel selection, maximizing the number of well-isolated single units in each target region.

#### Data Analysis

For all analyses, we used Python 3 with numpy (https://numpy.org/), scipy (https://docs.scipy.org/), matplotlib (https://matplotlib.org/), UMAP (https://github.com/lmcinnes/umap/blob/master/), sklearn (https://scikit-learn.org/stable/index.html)

#### Preprocessing single unit spiking

The spike times were rounded to 1,250 Hz. Then, we convolved the spike times of each unit with a 10 ms standard deviation gaussian kernel employing the convolve scipy function, which gave rise to a smoothed spiking activity^17^. For CA1 and CA3 recordings, recorded cells were divided into pyramidal cells, narrow interneurons, and wide interneurons based on the width of the waveform and the autocorrelogram. For RSC recordings, cells were divided into pyramidal cells and narrow interneurons. Only pyramidal cells in all regions were employed in the subspace analysis.

For Neuropixels 2.0 recordings on novel maze sessions we employed a total of 1171 CA1 pyramidal cells (97.6 ± 12.3 cells per session), 676 CA3 pyramidal cells (56.3 ± 12.7 cells per session), and 708 RSC pyramidal cells (59 ± 8.4 cells per session). For Neuropixels 2.0 recordings on two maze sessions we employed a total of 272 CA1 pyramidal cells (91.7 ± 15.7 cells per session), 74 CA3 pyramidal cells (24.7 ± 9.9 cells per session), and 119 RSC pyramidal cells (39.7 ± 12.4 cells per session). For SiNAPS recordings we employed a total of 191 CA1 pyramidal cells (63.7 ± 21.7 cells per session), 42 CA2 pyramidal cells (14 ± 4.6 cells per session), 41 CA3 pyramidal cells (13.7 ± 2.3 cells per session), and 56 DG pyramidal cells (18.7 ± 1.8 cells per session).

#### Partial canonical correlation analysis and mutual information

Canonical correlation analysis (CCA) has been used to characterize the communication subspace between distinct neural populations^17–19,100^. As a generalization to CCA, partial CCA (pCCA) is statistical technique to quantify the linear relationship between two sets of multivariables, after removing the influence of one or more covariate sets. It is useful when one wants to control for confounding effects in multivariate data analysis.

To conduct pCCA, we transformed the observations to meet the Gaussian assumption. All smoothed spike trains were z-scored by subtracting the mean firing rate and dividing the result by the standard deviation. Without loss of generality, given multivariate variables *X*, *Y* and *Z*, pCCA aims to seek a shared subspace between *X* and *Y* by finding sets of weights, *W_x_* and *W_y_*, that maximize the Pearson correlation coefficient ρ between the weighted *X* and weighted *Y* after removing the linear effects of *Z*. In essence, pCCA operates on the residuals of *X* and *Y* once *Z* has been regressed out. Let

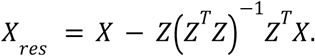

and

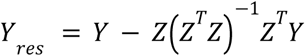

denote the residuals from *X* and *Y* (where Σ denotes the auto- or cross-covariance matrix for the respective variables labeled in the subscript) we computed a set of weight vectors such that

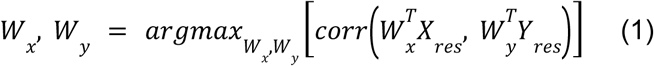

where 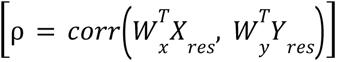 denotes the Pearson correlation coefficient. Mathematically, this is reduced to an expanded form of following equations

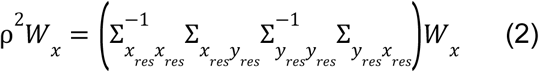

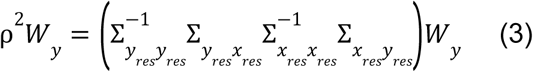

Solving the above equation is equivalent with the standard eigenvalue decomposition problem. By ranking their eigenvalues, we obtained from the largest to the smallest canonical correlation ρ. With computationally efficient eigenvalue decomposition, the *W*_*x*_ and *W*_*y*_ vectors were found as the eigenvectors of the scaled covariance matrices in the equations (2) and (3), respectively.

Note that there exist *M* pairs of *W*_*x*_ and *W*_*y*_ vectors that satisfy the previous equations, which are all the eigenvectors of the above-mentioned covariance matrices, where *M* = *min*(*N*_*x*_, *N*_*y*_), that is, the minimum number of neurons in either the *X* or *Y* population. Each *M* vector pair (referred here as pCCA components) defines one communication subspace (CS) pair. These components are ordered according to the canonical correlation coefficient (ρ) they produce, i.e., ρ(*CS*_1_) > ρ(*CS*_2_) > ρ(*CS*_3_) >… > ρ(*CS*_*M*_). We denote ρ_*max*_ = ρ_(_*CS*_1)_. Of note, because each CS component is one of the eigenvectors of the modified covariance matrix, they are orthogonal against each other, implying that the different CS dimensions for a given area are not correlated. A jupyter notebook demonstrating pCCA analysis is provided at: github/joaqgonzar

To generalize zero-lag to time-lagged pCCA, we computed the correlation between CS activities using the Pearson correlation coefficient *pearsonr* stats.scipy function. To find the optimal lag between pairs of areas, we temporally lagged one population by different times and computed pCCA to find the time-lag which maximizes the correlation ^17^.

In practice, we estimated the sample covariance matrices based on the temporally binned spike train data for *X* and *Y* (or *X*_*res*_ and *Y*_*res*_). When the dimensionality *N*_*x*_ *or* _*Ny*_ is very large, high-dimensional statistics implies that that a large sample size *N*≫*N*_*x*_, *N*_*y*_ is required to assure the covariance matrix estimate is accurate or stable ^101^. Empirically, we assured that the sample size was at least 1-2 orders of magnitude larger than the dimensionality of the variable. For instance, for a 100-dimensional variable, the minimum sample size requirement would be 50,000-500,000. For a given number of trials or recording duration, we could use a smaller number of bin size to increase the number of sample size *N*. Additionally, we varied the choice of bin size to assess the stability of subspace membership.

To understand how much information is transferred between two neuronal populations at different brain regions, we computed their mutual information (MI). From an information-theoretic perspective, MI measures how much uncertainty in one variable is reduced when knowing another variable ^102^. Assuming two multivariate variables are jointly Gaussian, we approximated MI using the following equation: 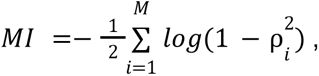 where 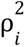 denotes the *i*-th squared canonical correlation value derived from pCCA.

#### Surrogate analysis

To understand how many significant CS pairs exist, we contrasted CS correlations through a surrogate analysis^17,19^. To generate a surrogate distribution to compare CS correlation values, we circularly shifted the spiking activity of one of the populations in time by random amounts (Extended Data Figure 3). For each surrogate data, we computed the pCCA and obtained the correlation coefficient for the first most correlated pair (CS_1_). The procedure was repeated 100 times for each session. We then assessed the significance by comparing the correlation coefficient for each CS pair against the null surrogate distribution. Only correlation values above the 95th percentile were deemed significant (P<0.05).

#### Rotation between subspaces

We quantified the rotation between two subspaces by computing its principal angles using the *subspace_angles* scipy function and then selecting the largest angle. Subspace angles, also known as principal angles, quantify the similarity between two subspaces. For two given subspaces, U and V, represented by orthonormal bases *Q*_*U*_ and *Q*_*V*_ respectively, the principal angles Θ are derived from the singular values of the matrix product 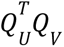 via Θ =arcos(σ), where σ_*i*_ are the singular values obtained from the singular value decomposition (SVD) of 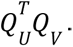

For the within subspace angle quantifications, we computed pCCA on a window of half of the recording session length and repeated this 100 times on different initial points. The maximum angle was computed for each iteration, and the median of the sine functions of these angles was computed. Then we applied arcsine to obtain the median angle. This allowed us to measure rotations within a subspace (same subspace in different windows) vs. rotations between subspaces. To compare across areas, we subsampled all areas to match the number of cells and repeated this procedure 100 times per session. The rotations reported correspond to the median maximum angle (applying the sine and then arcsine) over the 100 subsamples.

#### Projection of maze subspaces during sleep periods or different mazes

To project the subspaces computed during maze runs on different periods we stored the subspace weights *W* for all areas and then projected them as 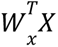 to yield a 1d signal for each set of weights. *X* corresponds to the population vector either during pre, post-maze or during the different maze run. We then computed Pearson’s correlation from the projected local activity to understand how correlations generalize across conditions.

#### Spatial information of pCCA weights

To quantify the spatial information contained in the pCCA weights we employed the following strategy: We first computed each cell’s place field by summing the number of spikes for each spatial bin (50 equally spaced bins across the length of the linear track) and dividing them by the bin time occupancy. Using a temporal bin size of 40 ms, we only employed temporal bins when the animal’s speed >2.5 cm/s. Both left and right runs were used. Place tuning curves were smoothed with a 12.5 cm gaussian kernel. We then extracted the place field for each cell as the bin showing the largest activity. Next, for each set of subspace weights we computed the average weight magnitude for each position along the track. To understand if subspace weight magnitudes were related to the gross position along the track (e.g., if weights for a given subspace were larger for cells with place fields at the end of the track), we divided position into 5 equally spaced bins and computed the average weight magnitude for each of these 5 bins. We then normalized these histograms by summing the activity across bins, yielding a probability distribution, and computed the entropy of each histogram. Thus, we obtained a measure for each subspace component quantifying how uniformly these weights were distributed in space. For control, we computed a surrogate distribution (100 repeats per subspace component) for each subspace component by circularly shifting each cell’s tuning curve by a random amount and computing the spatial entropy of its weights.

#### Variance explained by pCCA weights

We projected the subspace weights on the local activity as 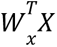 for each subspace dimension and then computed the variance for each projection to quantify the influence of cross-regional subspaces on local activity.

#### Dimensionality reduction via nonlinear UMAP method

Spike counts during maze running (speed > 2.5 cm/s) were binned into 40 ms bins and were then smoothed using a 500 ms wide Gaussian kernel and z-scored. The UMAP dimensionality reduction algorithm was then applied to the preprocessed data matrix. Each point in the low-dimensional manifold corresponds to the population activity at a single time bin. The UMAP code is available (https://github.com/lmcinnes/umap/blob/master/).

The UMAP hyperparameters were as follows: n_neighbors = 20, metric = ‘cosine’, output_metric = ‘euclidean’, learning_rate = 1.0, init = ‘spectral’, min_dist = 0.1, spread = 1.0, repulsion_strength = 1.0, negative_sample_rate = 5, target_metric = ‘categorical’, dens_lambda = 2.0, dens_frac = 0.3, dens_var_shift = 0.1.

To construct manifolds from subspace members, we only employed the spike trains from cells with absolute weights above the 75th percentile (|weight| > 0.75 quantile), while non-member neurons were defined as having absolute weights below the 25th percentile (|weight| < 0.25 quantile). Both manifolds were aligned to the full manifold using procrustes analysis (scipy).

#### Structure index for manifolds representation quantification

To quantify the degree to which the UMAP manifolds showed a meaningful structure for position and time, we computed the structure index^103^. The structure index (SI) quantifies how a given feature is structured along an arbitrary point cloud. To achieve this, we computed SI by first dividing the point cloud into several groups (bins) based on their feature values. These bins together cover the entire point cloud. For every pair of these bin groups, we then calculated an overlap score. This score represented the fraction of a bin group’s *k*-nearest neighbors (in the point cloud space) that belonged to the other bin group, based on a chosen distance measure like Euclidean distance. Computing the overlap score for each pair of bin groups (B_i_ and B_j_) yields an *n*×*n* adjacency matrix *M* whose entry (i,j) equals the overlap score between B_i_ and B_j_. Then, the structure index is defined as:

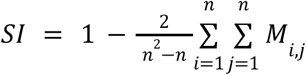

The SI statistic is ranged between 0 (random feature distribution) and 1 (maximally separated feature distribution).

#### Peer prediction analysis

We employed linear regression (*LinearRegression* sklearn function) to predict the activity of a given target neuron by either another subpopulation (upstream members/non-members, or local members/non-members, 75th/25th subspace weight threshold) of cells or by the spatial position of the animals. Spike counts during maze running (speed > 2.5 cm/s) were binned into 40 ms bins and were then smoothed using a 500 ms wide Gaussian kernel. 80% of the data was randomly divided to train the model and the remaining 20% was used for the test. The prediction score was averaged for each cell to yield a single value per session.

#### Place tuning curves and trajectory replay analysis

We built a Bayesian decoder during maze runs to predict the position of the animals based on spiking activity. Place tuning functions were first computed for each cell by summing the number of spikes for each spatial bin (2.5 cm) and dividing them by the bin time occupancy. Left and right runs were treated independently. Using a temporal bin size of 40 ms, we only employed temporal bins when the animal’s speed >5 cm/s. Place tuning curves were smoothed with a 12.5 cm gaussian kernel. A spatial modulation index (SMI) was derived from each cell normalized tuning curve (firing rate in each bin divided by the sum across bins) following SMI = (H_max_ - H)/H_max_, where H_max_ is the log(no.# of bins) and H denotes the Shannon Entropy of the normalized tuning curve. We quantified place field stability by computing the Pearson correlation (R) of the firing vectors between all trial combinations.

Only cells whose SMI>0.05 were included in the decoding analysis. Neurons across all areas and types that fit these criteria were pooled together. The median error of the decoded position was computed as a quality criterion and we only further analyzed sessions with the median decoding error <20 cm.

For replay analysis we identified candidate events during post-maze NREM sleep as the intervals where the population firing rate exceeded 2 standard deviations above the mean. Candidate events <50 ms or >500 ms were discarded. The spiking activity was then binned at 10 ms resolution and the posterior probability of position was computed for each bin in the candidate event. For each event we averaged the sequence score between the values obtained from the left run and right run decoders. To quantify replay quality, we computed the weighted correlation metric and generated a surrogate distribution of weighted correlations for each event by circularly shifting each cell’s place tuning (100 repeats). A sequence score was derived as:

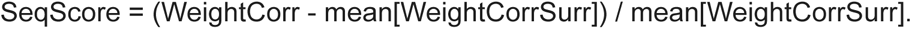

#### Cross-correlation analysis

We computed cross-correlograms with 1-ms resolution between two spike trains. The synaptic strength was estimated as the excess in causal spike transmission probability from that expected given *λ slow*^104^. Therefore, the normalized cross-correlogram (NCC) after n presynaptic spikes was defined as:

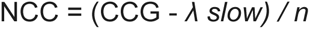

where λ slow is obtained by convolving the CCG with a partially hollowed gaussian kernel (with a standard deviation of 10 ms, with a hollow fraction of 60%).

#### Poisson spiking network simulation

We simulated spiking activity in a network of three neuronal populations (Populations X, Y, and Z), each composed of 5 neurons. Neurons emitted spikes as inhomogeneous Poisson processes with a baseline rate of 100 Hz, updated at a temporal resolution of 1 ms. The network contained two forms of interactions: cross-population modulatory connections (instantaneous excitatory effects between selected neurons in different populations) and local intra-population coupling (symmetric, reciprocal modulation within each population). The simulation proceeded in discrete timesteps of Δt=1 ms for a total duration of 100 s. At each timestep, the instantaneous probability of a spike for each neuron was computed as the sum of three components: 1) Baseline firing: Every neuron had a constant baseline firing probability corresponding to a Poisson firing rate of 100 Hz. 2) Cross-population modulation: Two dedicated modulatory pathways increased the instantaneous spike probability of specific neurons whenever a presynaptic “modulator” neuron spiked in the same timestep: Z→X,Y modulation: A single neuron in Population Z (Z₂) acted as a modulatory source for one neuron in Population X (X₀) and one neuron in Population Y (Y₀). If Z₂ spiked, it instantly boosted the spike probability of its targets by an additional factor (see Extended Data Fig. 4). X→Y modulation: If the neuron X₀ spiked, it instantly boosted the spike probability of Y₀ by a given factor. All cross-population effects were *instantaneous*: if a modulator neuron spiked at time t, its target neurons received the modulation within the same timestep and had an elevated spike probability in that same millisecond. 3) Local intra-population coupling. When enabled, each population had a symmetric connectivity local matrix describing the strength of reciprocal connections between all neuron pairs. If any neuron in a population spiked at timestep t, all other neurons in the same population received an instantaneous additive input. This additional input increased their spike probability within the same timestep, allowing synchronous co-activation.

Four simulation conditions were designed by systematically varying the strength of (i) modulatory inputs X to Y (tenfold increase in spiking probability), (ii) modulation from Population Z to X and Y (tenfold increase in spiking probability), (iii) simultaneous modulation from Z to X and Y (tenfold increase in spiking probability) and also an independent modulation from X to Y (fivefold increase in spiking probability), and (iv) local intra-population connectivity (tenfold increase in spiking probability) on top of the scenario described in (iii). Note that we allowed different base firing rates for the simulation shown in panel e of Extended Data Figure 4.

#### Statistics

We employed non-parametric statistics throughout the manuscript. For paired data we employed Wilcoxon’s signed-rank test, while we used the ranksum test for unpaired data or the Kolmogorov-Smirnov (KS) test to compare two distributions. When appropriate we constructed surrogate distributions from the recorded data. We set P<0.05 as the threshold for significance.

## Data availability

The experimental data is available upon request to the corresponding authors.

## Code availability

All original code developed for this project is available at: https://github.com/joaqgonzar/pCCA-Hippocampus Two Jupyter notebooks are provided demonstrating pCCA use with simulated and real data.

## Acknowledgements

We thank Horacio Rotstein, Gergely Komlosi, and Adriano Tort for providing insightful comments on our manuscript. We thank Elisa Chinigò for providing feedback on the spiking network model. The work was supported by grants DA056394 (Z.S.C., B.G.), R01MH122391 and U19NS107616 (G.B.) from the US National Institutes of Health. J.G. was supported by the Uruguayan Sistema Nacional de Investigadores (SNI) and a Ben Barres Spotlight Award from eLife.

## Author contributions

M.V., R.S., and G.B. designed the experiments. M.V., N.S., D.A., and N.N., performed the experiments and data curation. J.G. analyzed the data with the supervision of Z.S.C. J.G., Z.S.C., and G.B. wrote the original manuscript with inputs from all authors.

## Supplementary Materials

**Extended Data Figure 1.**
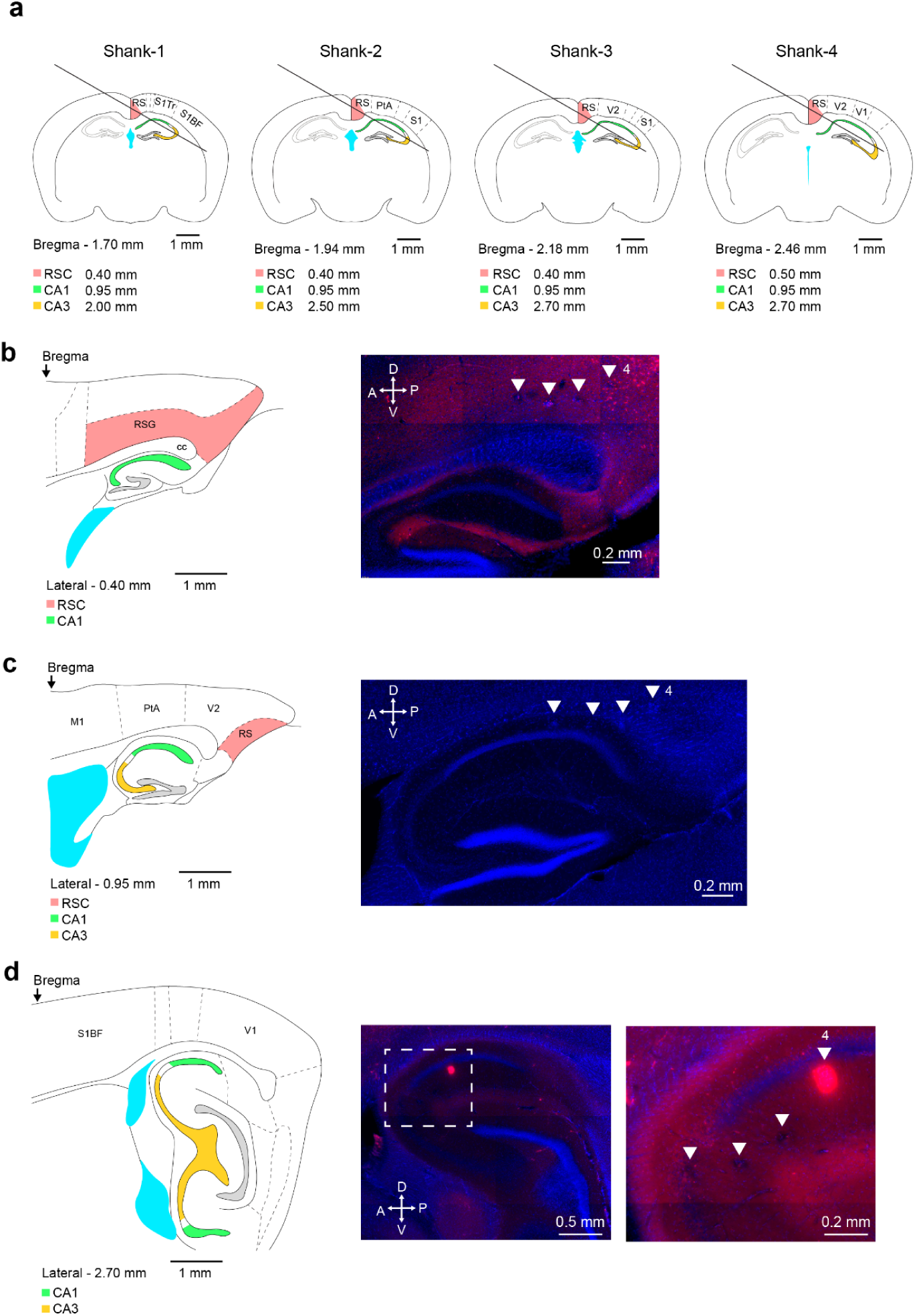
Brain atlas schematics and histological reconstruction of probe tracks. **a** Schematics of coronal sections show the location of each shank of the NP2.0 probe relative to Bregma. Red, green and yellow highlights granular RSC, CA1 and CA3, respectively. Note that shank-4 reaches CA3 at midline – 2.7 mm compared to shank-1 that penetrates CA3 at midline – 2.00 mm. **b-d** Brain atlas schematic of sagittal sections at midline -0.4 mm. **b** -0.95 mm (**c**), and -2.70 mm **d** and corresponding histological sections (right) stained with DAPI (blue) and DiI (red). White triangles show the location of the four shanks. DiI was applied on the tip of shank-4 only. Note the strong signal in the red channel in **d** (left section). Right section shows the zoomed-in version of the right dashed box from the left.

**Extended Data Figure 2.**
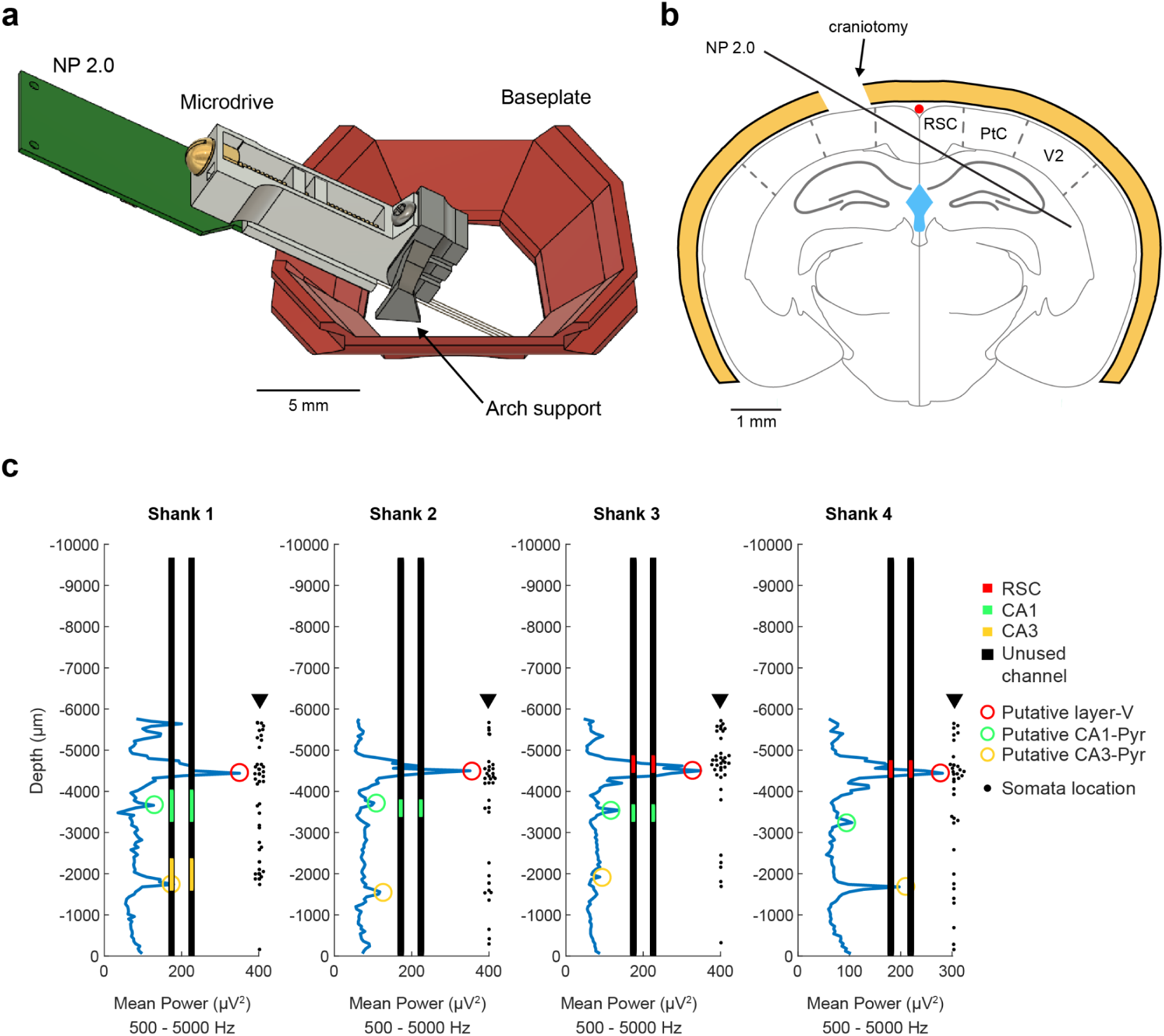
Surgical procedure and layer identification. **a)** 3D rendering of plastic baseplate (red), metal microdrive and NP2.0 probe. The baseplate contains grooves on both sides, allowing head fixation of chronically implanted mice. **b)** Coronal section 2 mm posterior from Bregma, showing the location of the craniotomy and the brain regions that the NP2.0 probe can record from. **c**) 4-shank Neuropixels (NP 2.0) recording. Recorded channels spanned 5.7 mm on each shank (60 µm spacing between active channels in one row). Blue line is the multi-unit spectral power (500 Hz to 5 kHz) distribution as a function of depth). Circles show the maximum peaks corresponding to cellular layers (red – putative layer 5 of RSC, green – putative CA1 pyramidal layer, yellow – putative CA3 pyramidal layer). Black rectangles show the recording channels of NP2.0. Colored squares (red, green and yellow) show the selected channels for simultaneous recording. Black dots show the somata location of the recorded units. Note high-density of somata around the cellular layers. Note also that simultaneous RSC recordings were made from the same span as in CA1 and CA3 from layers 5/6. For simplicity, were refer to “superficial” and “deep sublayers” as the relative position of the neurons in layers 5/6.

**Extended Data Figure 3.**
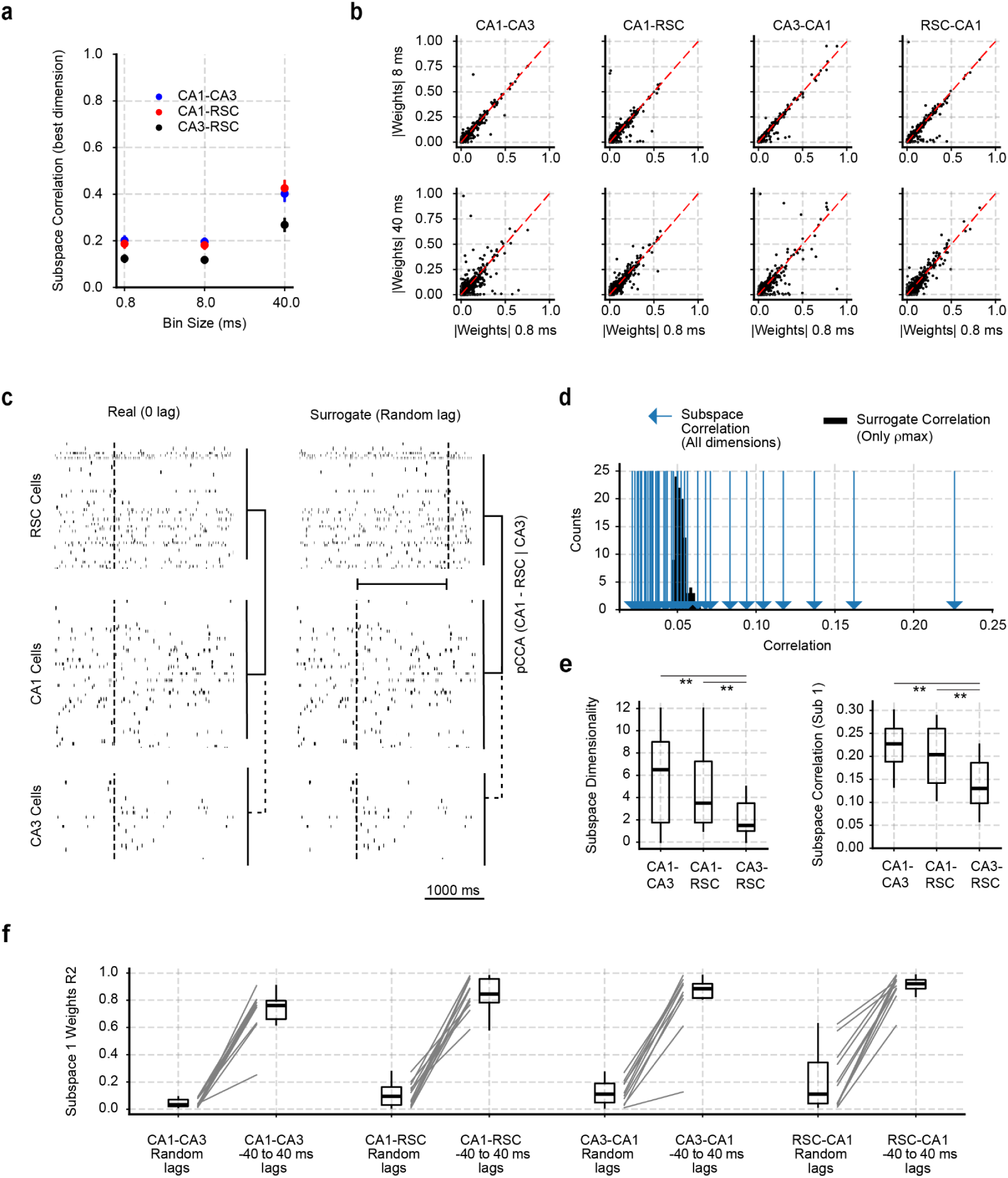
Subspace timescale and dimensionality. **a** Subspace correlation (for the best subspace) as a function of the temporal bin size employed in the analysis. **b** Correlation between the subspace weights (best subspace) computed at different time bins. Note cell memberships are consistent irrespective of the time bin used. Thus, we chose the smallest time bin to gain statistical power in the analysis. **c** Schematic of the surrogate test for subspace significance. One of the populations is circularly shifted with respect to the other and pCCA is computed for 100 different iterations for each session. Only the best subspace in each surrogate is retained to make the surrogate distribution. **d** Distribution of surrogate pCCA correlation (black). Real values are superimposed by the blue lines. Only subspaces with P<0.05 are considered significant. **e** Subspace dimensionality and correlation for the NP 2.0 experiments. **f** Correlations (R^2^) between the best subspace computed at lag 0 (between the two populations), compared to either a random lag (100 random lags per session), or to a lag within a -40 to 40 ms window. Note that subspace weights computed for random lags retain almost no information about the real cell memberships. P values were obtained through a Wilcoxon signed-rank test.

**Extended Data Figure 4.**
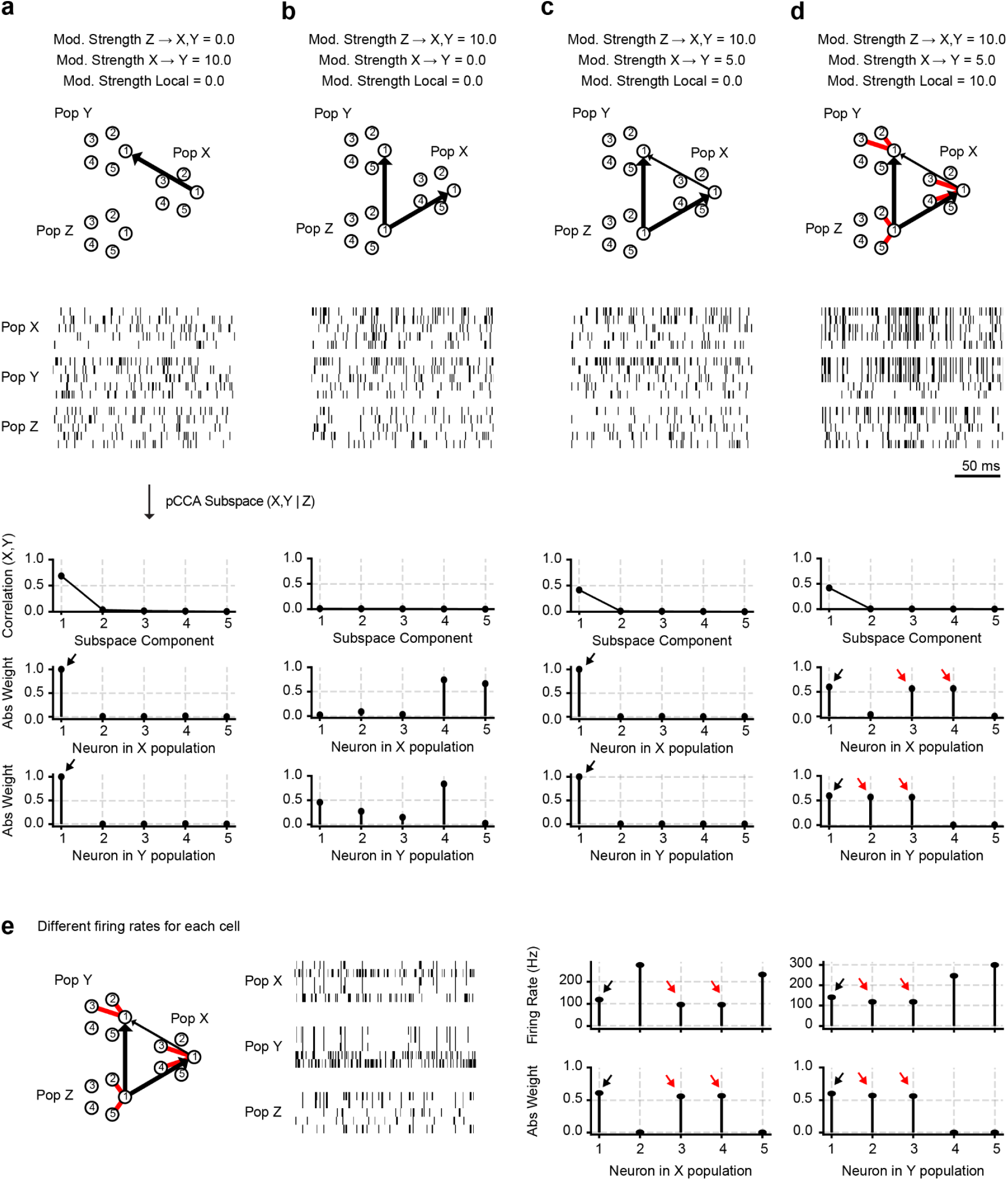
Recovering genuine pairwise subspace interactions through pCCA. Simulated Poisson spiking activity in three interconnected populations (X, Y, Z) under different modulation conditions. Top: Connectivity diagrams show modulation strength from population Z to X/Y and from X to Y. Mod strengths on top indicate fold change from baseline firing rate (100 Hz). Middle: Example raster plots of spiking activity for each condition. Bottom: Subspace analysis results: the first row shows the correlation strength captured by successive subspace components; the second and third rows display absolute subspace weights for individual neurons in populations X and Y. **a** Strong modulation from X → Y produced a dominant single subspace component with selective neuron weights (arrows). **b** Strong modulation from Z → X/Y drove synchronized inputs to both X and Y, by conditioning by Z activity, pCCA was able to overcome this third-party modulation. Note that subspace correlation dropped to 0. **c** Combined modulation from Z → X/Y and moderate X → Y coupling. pCCA preserves a dominant low-dimensional subspace with selective membership provided by the X → Y coupling. **d** Adding strong local modulation generated a richer structure with multiple neurons (red arrows) contributing to the subspace. Note that even though local dynamics might be strong/dominant, pCCA is still effective in recovering the genuine long range interactions. **e** Same example as in **d** but having different base firing rates for each cell in each area. We specifically made cells connected across regions to have lower firing rates compared to non-connected cells, thus showing that pCCA is still able to identify the connected group despite their lower number of spikes produced.

**Extended Data Figure 5.**
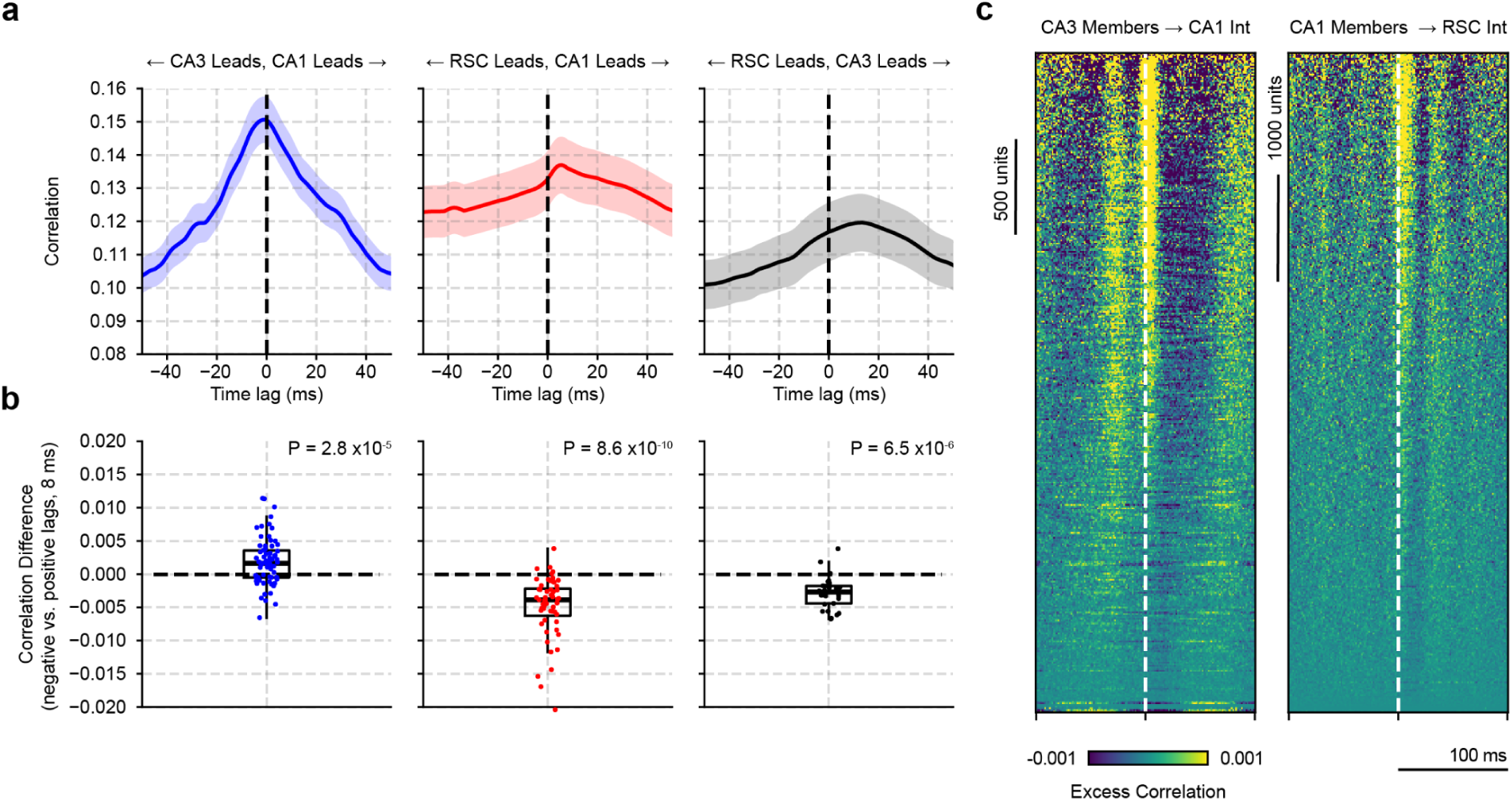
Subspace directionality. **a** Temporal lag analysis. One of the populations was lagged with respect to the other and pCCA was quantified for each lag. CA3 leads CA1 (left), RSC lagged CA1 (middle), whereas CA3 leads RSC. **b** Quantification of asymmetry in temporal correlations for each pair, confirming significant lead–lag structure for CA3→CA1 and CA1→RSC and CA3→RSC (p-values shown). Each dot shows a single subspace correlation difference. Data were pooled across animals and sessions. P values were obtained through a Wilcoxon signed-rank test. **c** Spike-triggered population activity for CA3 subspace members (cells above the 75th weight quantile in the best subspace) and CA1 interneurons (left) and CA1 members (same 75th quantile threshold) and RSC interneurons (right) revealed temporally precise activation patterns aligned with inferred lead–lag directions.

**Extended Data Figure 6.**
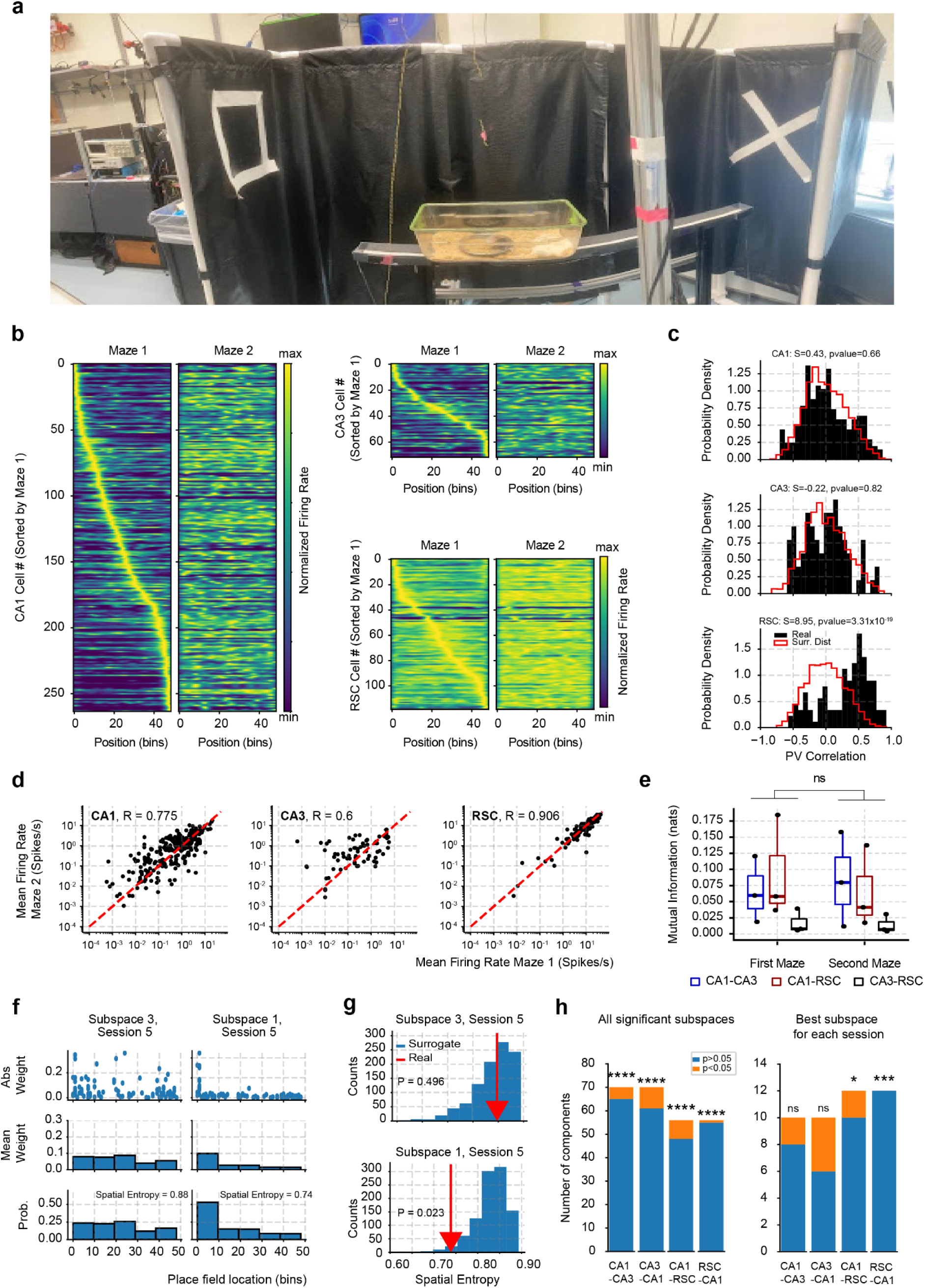
Remapping in the two-maze experiments and spatial information of subspace weights. **a** Photographs of the two mazes in two room configurations **b** Normalized firing rate for all recorded cells as a function of location on the linear track. **c** Population vector (PV) correlations between the first and second maze tuning curves. Black histogram shows the real distribution of PV correlations pooling all analyzed cells. Red histogram shows a surrogate distribution constructed by randomly circularly shifting the second maze tuning curve (100 times for each cell). P values were obtained through a Wilcoxon ranksum test. **d** Firing rate correlations in both mazes. Note the high correlation of log firing rates, despite remapping of place fields. **e** Mutual information between pairs of brain regions (CA1–CA3, CA1–RSC, CA3–RSC) in Maze 1 and Maze 2. Mutual information was computed from the subspace correlation (see Methods)**. f** Example of the spatial information carried by subspace weights. Two significant subspaces are shown, one lacking any spatial selectivity in the subspace weights and other showing larger weights for cells firing at the beginning of the track. **g** Surrogate testing conducted by shifting place field identity across cells shown in panel **b**. The amount of spatial information is measured by the spatial entropy of the subspace magnitude x place field location histograms. **h** Number of subspaces showing spatial organization.

**Extended Data Figure 7.**
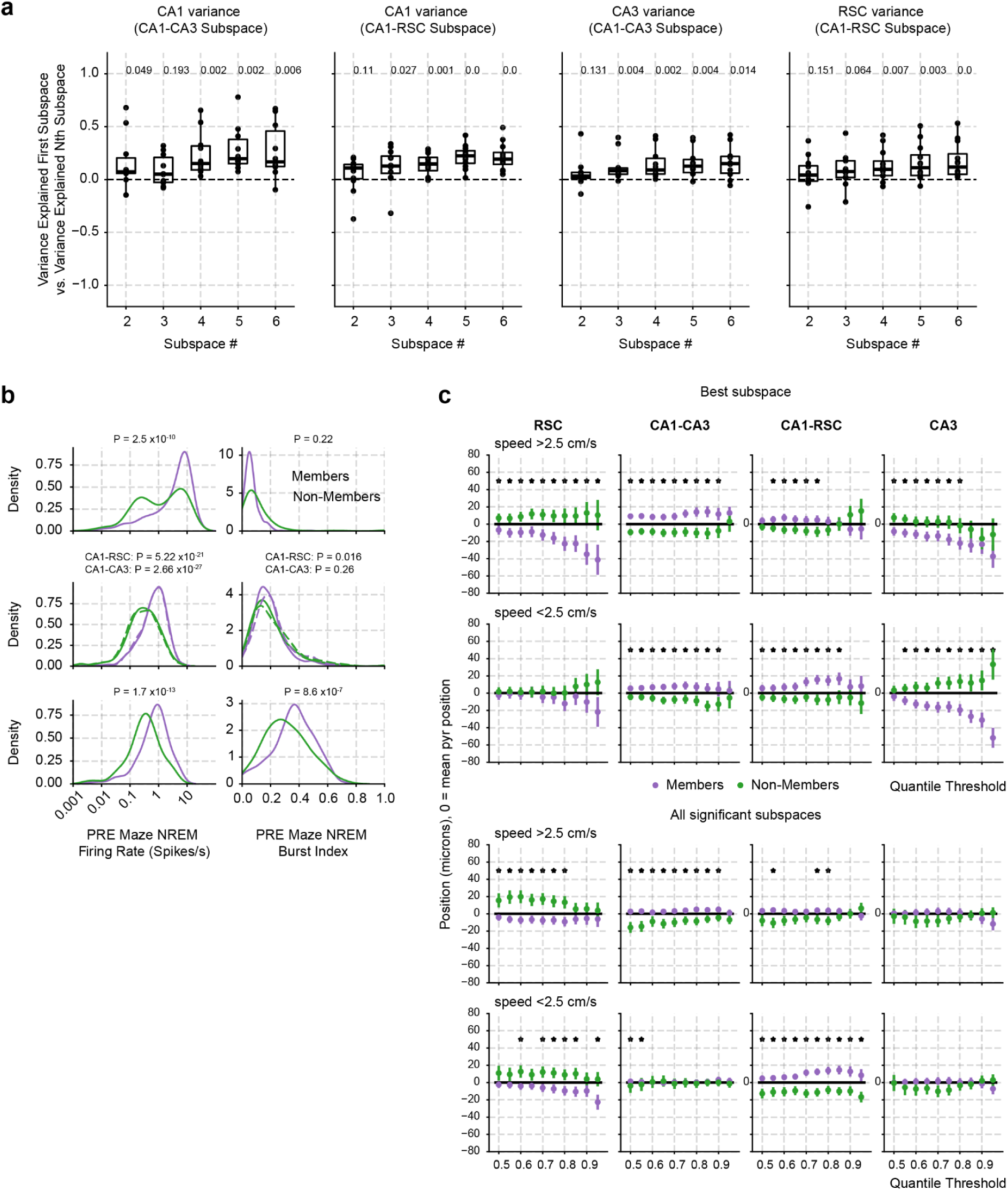
Subspace members and non-members show distinct physiological and anatomical properties. **a** Variance explained by the difference between the first subspace and the remaining subspaces. P values were obtained through a Wilcoxon signed-rank test. **b** Firing rate distributions for members of all significant subspaces. **c** Influence of quantile threshold, running speed, and the number of subspace dimensions employed. P values were obtained through a Wilcoxon ranksum test.

**Extended Data Figure 8.**
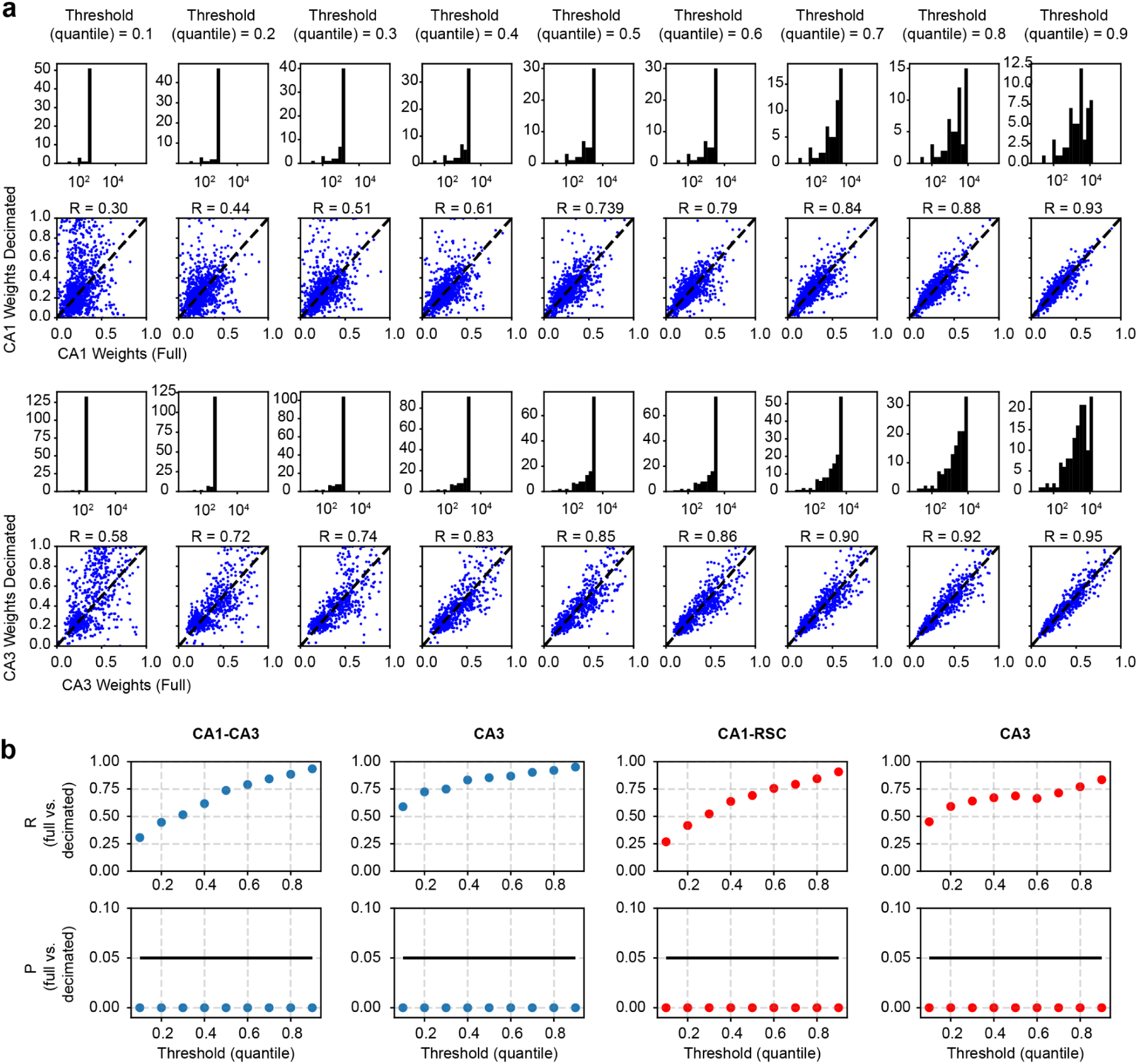
Firing rate controls for subspace memberships. **a** Random subsampling of spikes to match all neurons to the *i*-th population quantile. No spikes were removed if the total number of spikes for a given neuron was below such a threshold. Top histograms: Distribution of the number of spikes for all neurons. Bottom scatter plots: Maximum absolute subspace weight for each cell before and after subsampling (all subspaces were considered for this analysis). **b** Correlation coefficients (top row) and P values (bottom row) between the subsampled and non-subsampled subspace weights.

**Extended Data Figure 9.**
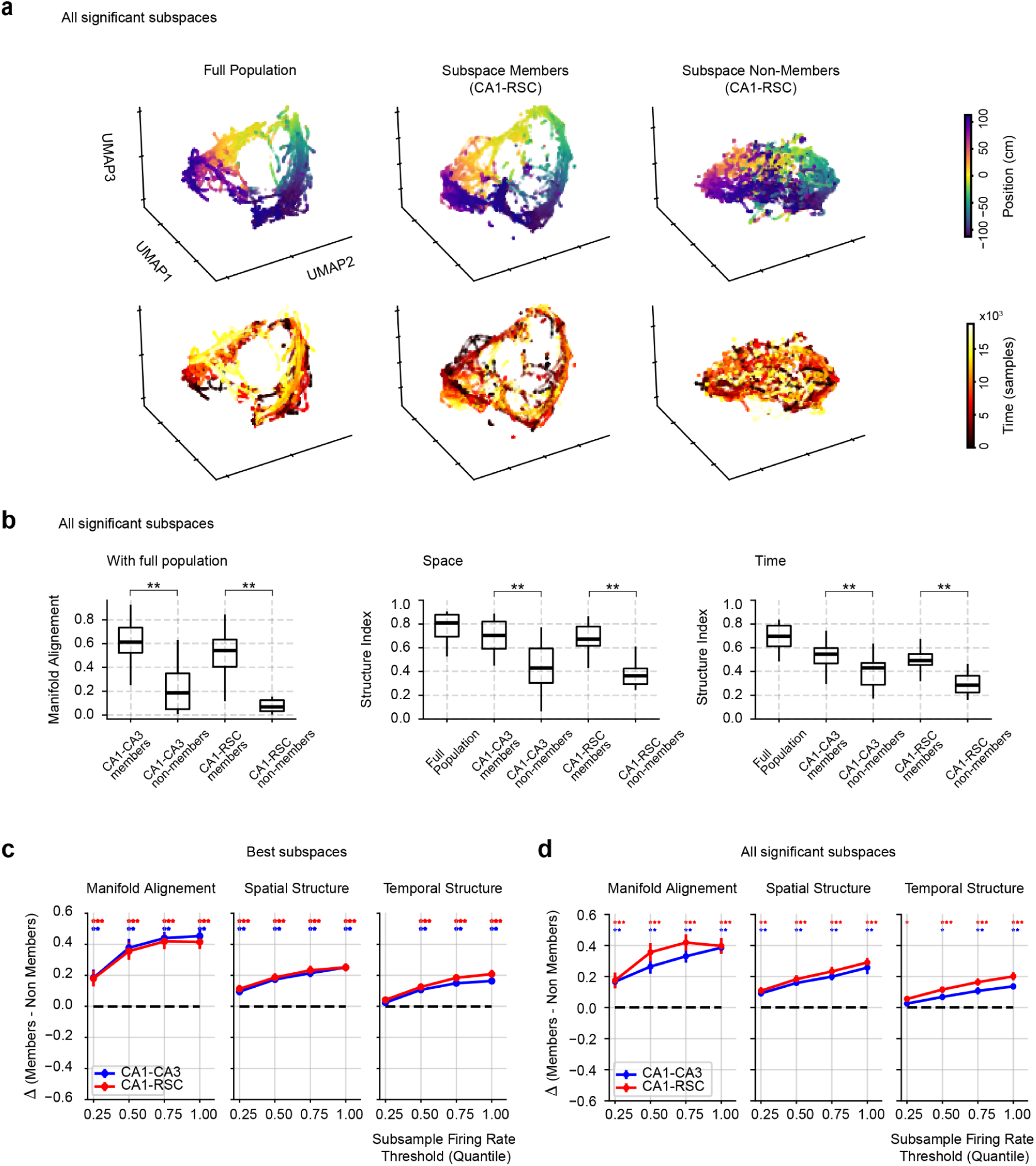
Quantifying UMAP values and firing rate control. **a** UMAP manifold computed for members and non-members of all significant subspaces (0.9 and 0.1 thresholds for each subspace). **b** Left: Manifold alignment to full population manifold. Middle: Spatial structure index for UMAP embeddings produced from subspace members across all significant subspaces. The structure index quantifies the extent to which the distribution of a given feature (position or trial) deviates from a uniform distribution in the UMAP point cloud. Right: Temporal structure index for UMAP embeddings. **c** Subsampling analysis and UMAP quantification as a function of firing rate quantiles. Members were computed for the best subspace for each session (0.75 and 0.25 thresholds). **d** Subsampling analysis and UMAP quantification as a function of firing rate quantiles. Members were computed for all significant subspaces for each session (0.75 and 0.25 thresholds). P values were obtained through a Wilcoxon signed-rank test.

**Extended Data Figure 10.**
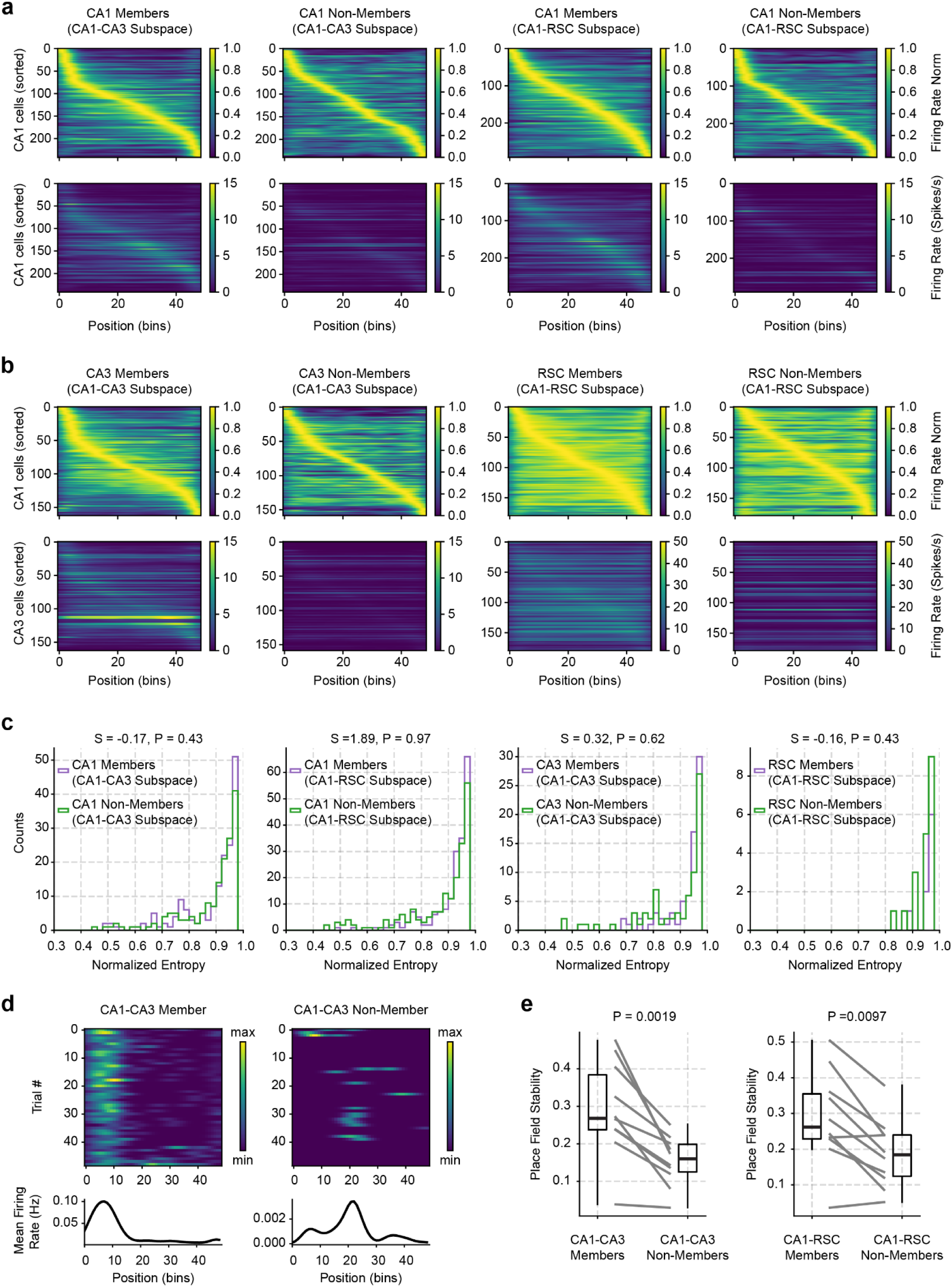
Place fields for subspace members and non-members. **a,b** Normalized (top) and absolute (bottom) firing rates (Hz) for all subspace member and non-member cells. All significant subspaces were employed for this analysis. 0.75 and 0.25 quantiles were chosen as thresholds for member and non-member definitions, respectively. All cells were pooled across sessions. **c** Normalized entropy for each cell’s tuning curve. Statistics shown on the top of each panel were computed from the rank-sum test. Individual place field features of subspace member and non-member neurons did not differ, in contrast to their difference in temporal organization (Fig. 4 and Extended Data Figure 9). P values were obtained through a Wilcoxon ranksum test. **d** Top: Firing rate as a function of position and trial number. Bottom: Average firing rate across all trials. **e** Firing rate correlation (R) across all trial combinations averaged for each session for members and non-members. P values were obtained through a Wilcoxon signed-rank test.

**Extended Data Figure 11.**
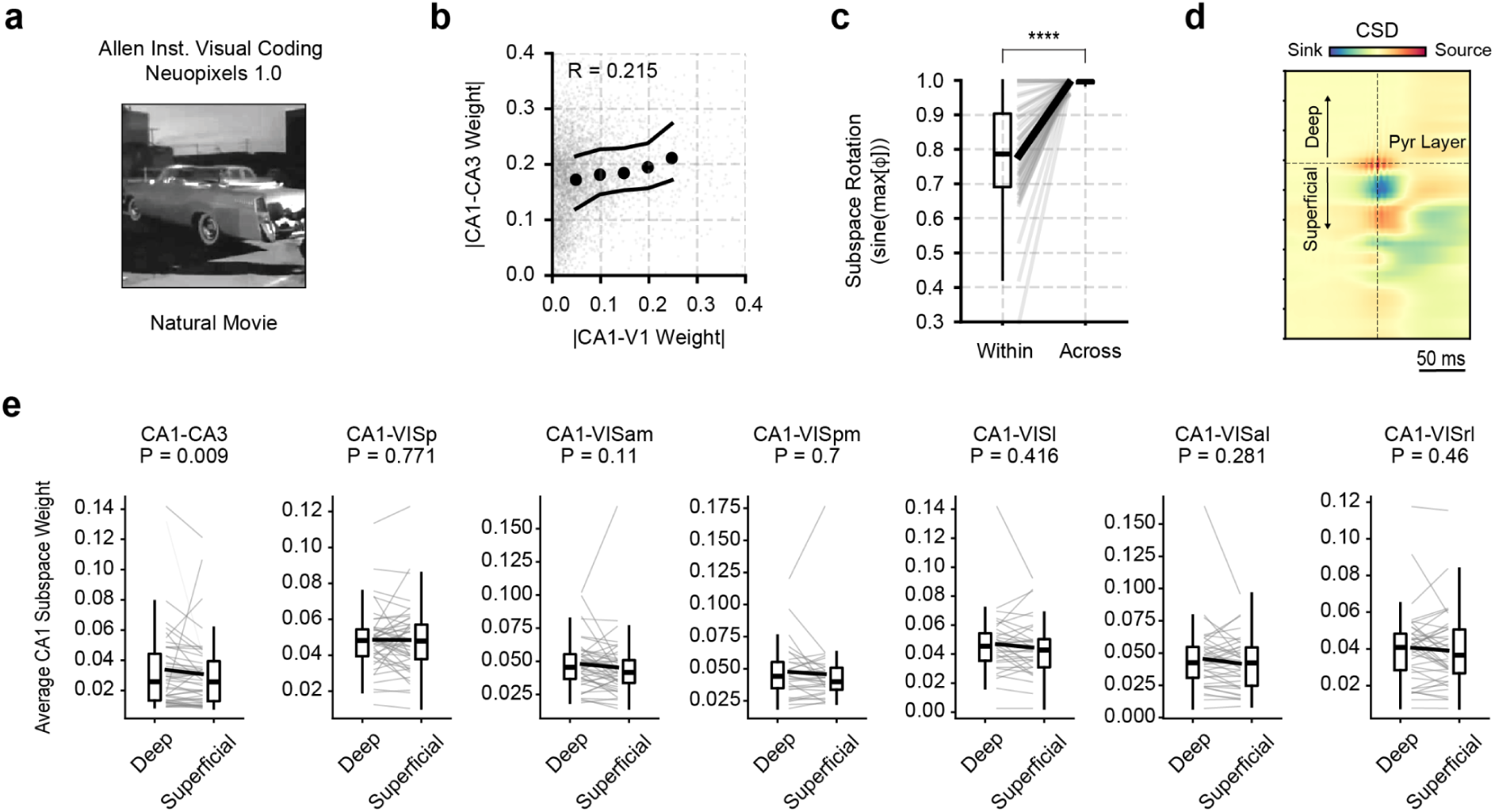
Deep sublayer CA1 neurons preferentially contribute to hippocampal–but not visual–subspaces during movie viewing. **a** Schematic of the Allen Institute Visual Coding paradigm. Mice passively viewed natural movies while neuronal activity was recorded with Neuropixels 1.0 probes spanning the hippocampus and visual cortex. **b** Significant correlation between CA1 subspace weights for CA1–V1 and CA1–CA3 pairs (R = 0.215), suggesting partial overlap in CA1 neuron participation across hippocampal and visual interactions. **c** Subspace rotation analysis between CA1–CA3 and CA1–V1 subspaces (****P < 0.0001, paired test). **d** Current source density (CSD) profile aligned to ripple peak. **e** Average CA1 subspace weights split by anatomical depth (deep vs. superficial layers) across CA1–CA3 and CA1–visual cortical area pairs. Deep-layer CA1 neurons contribute more strongly to CA1–CA3 subspaces (P = 0.009), but not significantly to any CA1–visual subspace (all P > 0.1), suggesting that hippocampal–visual coupling was drawn from a broader subset of CA1 cells. P values were obtained through a Wilcoxon signed-rank test.

**Extended Data Figure 12.**
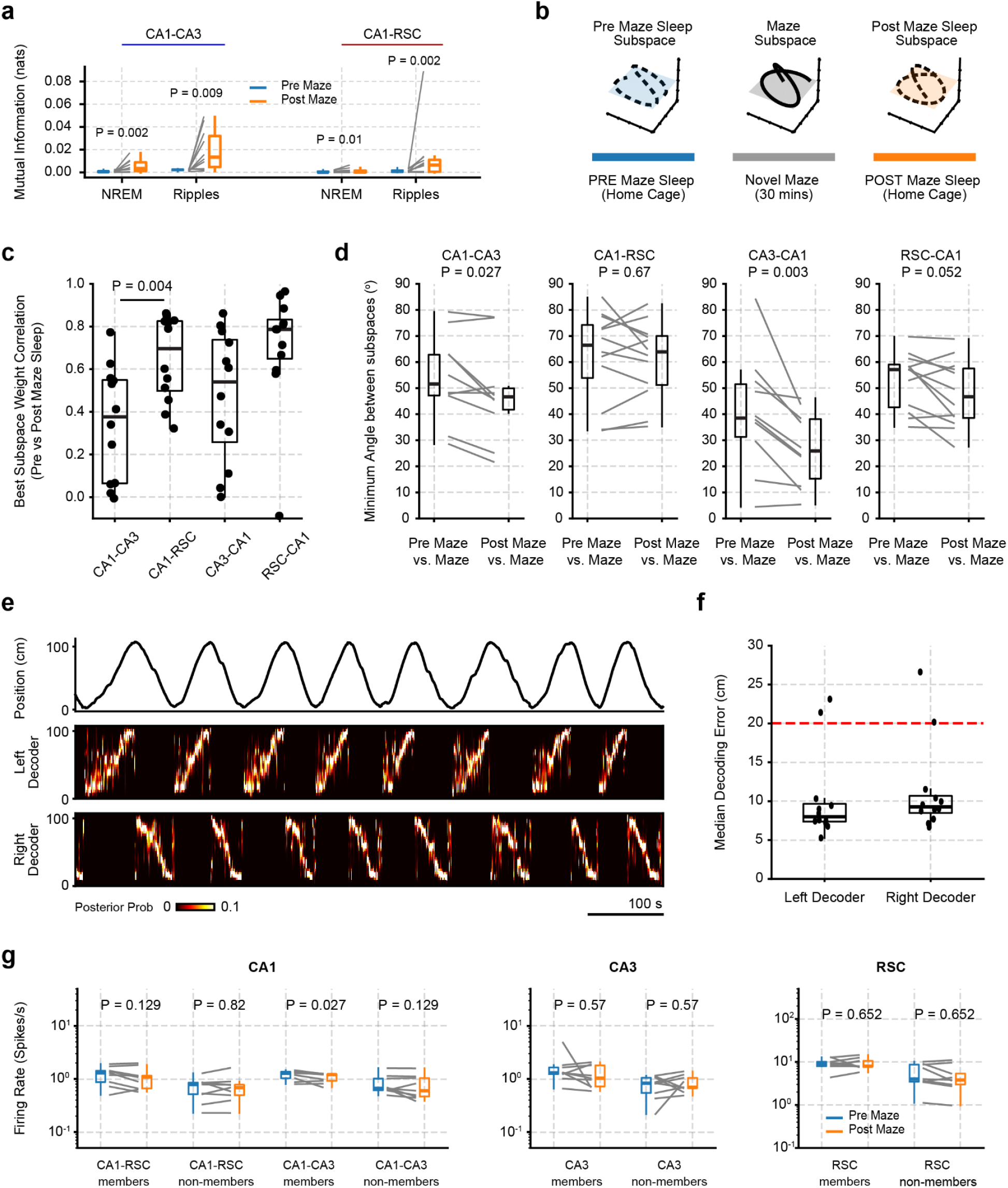
Firing rate changes during pre-experience and post-experience sleep. **a** Subspace mutual information (all significant subspaces) between maze behavior versus NREM and SPW-R events, quantified as mutual information **b** Schematic showing the independent computation of each subspace during pre-maze, maze, and post-maze periods. **c** Subspace |weight| correlation between pre and post maze periods. Only the best subspace was used for this analysis. **d** Minimum principal angle between pre/post and maze subspaces. All significant subspaces were employed for this analysis. **e** Illustrations of Bayesian decoder trained to predict the animal’s position during maze runs. A separate decoder was employed for left and right runs. **f** Median absolute error for all sessions (median[abs(Decoded Position - Real position))]. A threshold of 20 cm (red line) was employed as an inclusion criterion for the analysis. **g** Firing rates across all cells for a given group were averaged for each session. P values were obtained through a Wilcoxon signed-rank test.

**Extended Data Figure 13.**
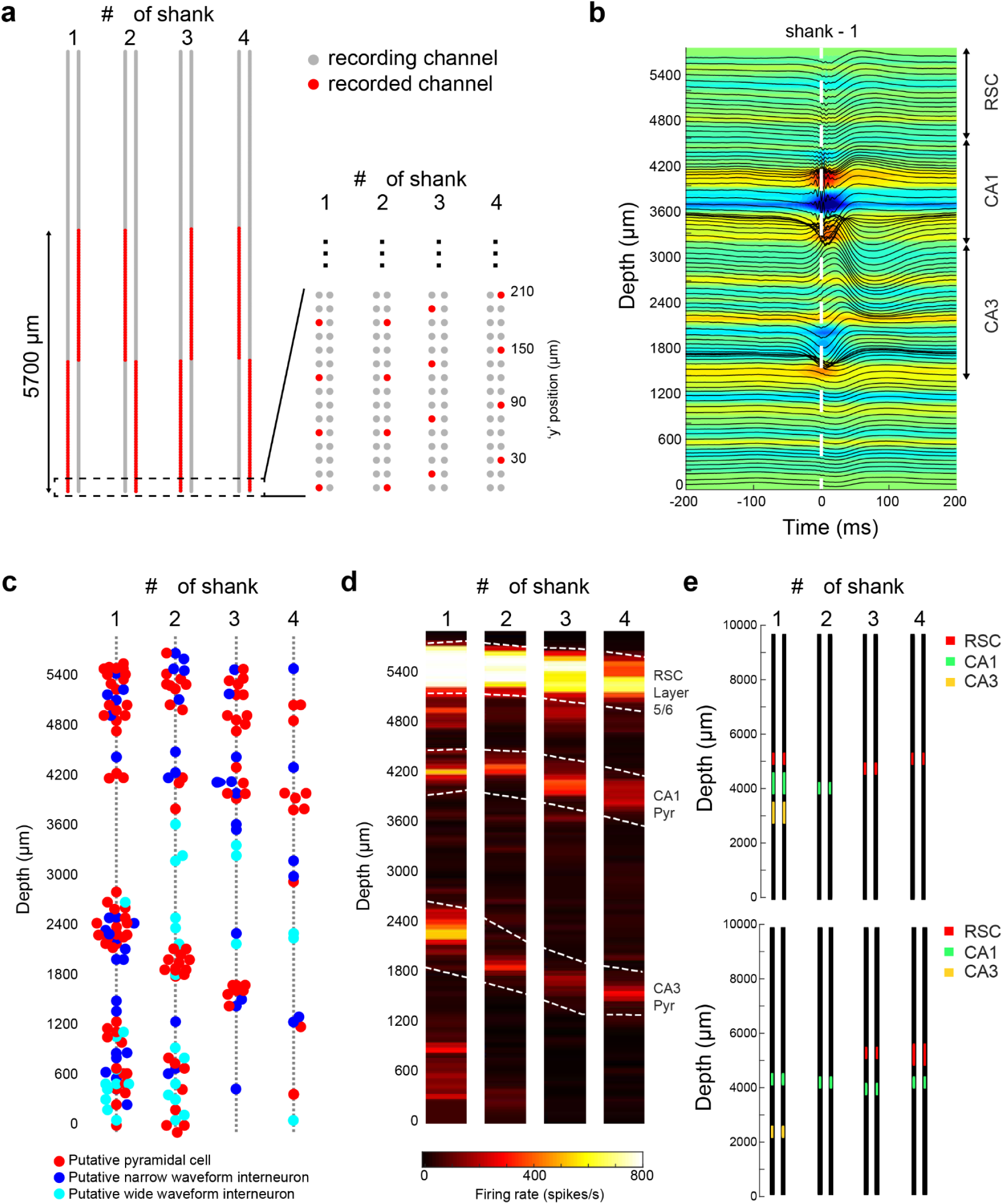
Identification of brain regions of interest using Neuropixels 2.0 probe. **a)** Neuronal activity was recorded across 4 shanks spanning 5760 µm by 750 µm (every 4^th^ channel was selected on each shank to create a linear probe profile). Red channels show the selected channels on each shank. Right: zoomed in on the probe tip. Grey circles show available recording channels, red circles show selected channels. **b)** Average current source density (CSD, color) map and superimposed LFP traces of SPW-R events (± 200 ms). Note negative sharp waves and sinks (blue) in the stratum radiatum of CA1. **c)** Probe layout is shown with the putative location of recorded neuron somata (n = 97 putative pyramidal cells, 56 narrow interneurons and 58 wide interneurons). Single units were clustered in the cellular layers of cortex and hippocampus (0 µm represents the tip of the probe). **d)** Population firing rate measured from the high-pass filtered signal (500 Hz) crossings of a −50 µV threshold, in a 10-s interval. **e)** Example channel maps selected for two different sessions. 8 blocks of 48 channels were distributed across RSC (red), CA1 (green) and CA3 (yellow).

**Supplementary Table 1.** Mazes used for behavioral experiments.

| Maze name | Length (cm) | Width (cm) | Material | Coating |
| --- | --- | --- | --- | --- |
| Pre-training | 110 | 5 | Plastic (McMaster 85065K55) | Silicone (Silicone Heat Resistant Mat) |
| Maze-1 | 110 | 5 | Plastic (McMaster 85065K55) | Gaffer Power tape (white) |
| Maze-2 | 110 | 5 | Fiberglass (McMaster 8529K41) | Gaffer Power tape (white) |
| Maze-3 | 100 | 7 | Wood | White paint |

## References

1. Semedo, J. D., Zandvakili, A., Machens, C. K., Yu, B. M. & Kohn, A. Cortical Areas Interact through a Communication Subspace. Neuron 102, 249–259.e4 (2019).

2. Squire, L. R., Genzel, L., Wixted, J. T. & Morris, R. G. Memory consolidation. Cold Spring Harb. Perspect. Biol. 7, a021766 (2015).

3. Buzsáki, G. Two-stage model of memory trace formation: a role for ‘noisy’ brain states. Neuroscience 31, 551–570 (1989).

4. Alvarez, P. & Squire, L. R. Memory consolidation and the medial temporal lobe: a simple network model. Proc Natl Acad Sci U S A 91, 7041–7045 (1994).

5. Marr, D. Simple memory: a theory for archicortex. Philos Trans R Soc Lond B Biol Sci 262, 23–81 (1971).

6. Squire, L. R. Memory and the hippocampus: a synthesis from findings with rats, monkeys, and humans. Psychol Rev 99, 195–231 (1992).

7. Lavenex, P. & Amaral, D. G. Hippocampal-neocortical interaction: a hierarchy of associativity. Hippocampus 10, 420–430 (2000).

8. O’Keefe, J. & Nadel, L. The Hippocampus as a Cognitive Map. (Oxford University Press, USA, 1978).

9. Chaudhuri, R., Gerçek, B., Pandey, B., Peyrache, A. & Fiete, I. The intrinsic attractor manifold and population dynamics of a canonical cognitive circuit across waking and sleep. Nat Neurosci 22, 1512–1520 (2019).

10. Sussillo, D., Churchland, M. M., Kaufman, M. T. & Shenoy, K. V. A neural network that finds a naturalistic solution for the production of muscle activity. Nat Neurosci 18, 1025–1033 (2015).

11. Remington, E. D., Egger, S. W., Narain, D., Wang, J. & Jazayeri, M. A Dynamical Systems Perspective on Flexible Motor Timing. Trends Cogn Sci 22, 938–952 (2018).

12. Wang, J., Narain, D., Hosseini, E. A. & Jazayeri, M. Flexible timing by temporal scaling of cortical responses. Nat Neurosci 21, 102–110 (2018).

13. Sadtler, P. T. et al. Neural constraints on learning. Nature 512, 423–426 (2014).

14. Oby, E. R. et al. Dynamical constraints on neural population activity. Nat Neurosci 28, 383–393 (2025).

15. Kaufman, M. T., Churchland, M. M., Ryu, S. I. & Shenoy, K. V. Cortical activity in the null space: permitting preparation without movement. Nat Neurosci 17, 440–448 (2014).

16. Kohn, A. et al. Principles of Corticocortical Communication: Proposed Schemes and Design Considerations. Trends Neurosci 43, 725–737 (2020).

17. Gonzalez, J., Torterolo, P., Bolding, K. A. & Tort, A. B. L. Communication subspace dynamics of the canonical olfactory pathway. iScience 27, 111275 (2024).

18. Kim, J., Joshi, A., Frank, L. & Ganguly, K. Cortical-hippocampal coupling during manifold exploration in motor cortex. Nature 613, 103–110 (2023).

19. Veuthey, T. L., Derosier, K., Kondapavulur, S. & Ganguly, K. Single-trial cross-area neural population dynamics during long-term skill learning. Nat Commun 11, 4057 (2020).

20. Ebrahimi, S. et al. Emergent reliability in sensory cortical coding and inter-area communication. Nature 605, 713–721 (2022).

21. Jun, J. J. et al. Fully integrated silicon probes for high-density recording of neural activity. Nature 551, 232–236 (2017).

22. Steinmetz, N. A. et al. Neuropixels 2.0: A miniaturized high-density probe for stable, long-term brain recordings. Science 372, (2021).

23. Steinmetz, N. A., Zatka-Haas, P., Carandini, M. & Harris, K. D. Distributed coding of choice, action and engagement across the mouse brain. Nature 576, 266–273 (2019).

24. Buzsáki, G. Hippocampal sharp wave-ripple: A cognitive biomarker for episodic memory and planning. Hippocampus 25, 1073–1188 (2015).

25. Muller, R. U. & Kubie, J. L. The effects of changes in the environment on the spatial firing of hippocampal complex-spike cells. J Neurosci 7, 1951–1968 (1987).

26. Leutgeb, S. et al. Independent codes for spatial and episodic memory in hippocampal neuronal ensembles. Science 309, 619–623 (2005).

27. Mizuseki, K., Diba, K., Pastalkova, E. & Buzsáki, G. Hippocampal CA1 pyramidal cells form functionally distinct sublayers. Nat Neurosci 14, 1174–1181 (2011).

28. Nitzan, N. et al. Propagation of hippocampal ripples to the neocortex by way of a subiculum-retrosplenial pathway. Nat Commun 11, 1947 (2020).

29. Leland, M., John, H. & James, M. UMAP: Uniform Manifold Approximation and Projection for Dimension Reduction. arXiv (2020) doi:10.48550/arXiv.1802.03426.

30. Mizuseki, K., Royer, S., Diba, K. & Buzsáki, G. Activity dynamics and behavioral correlates of CA3 and CA1 hippocampal pyramidal neurons. Hippocampus 22, 1659–1680 (2012).

31. Harris, K. D., Csicsvari, J., Hirase, H., Dragoi, G. & Buzsáki, G. Organization of cell assemblies in the hippocampus. Nature 424, 552–556 (2003).

32. Grosmark, A. D. & Buzsáki, G. Diversity in neural firing dynamics supports both rigid and learned hippocampal sequences. Science 351, 1440–1443 (2016).

33. Skaggs, W. E. & McNaughton, B. L. Replay of neuronal firing sequences in rat hippocampus during sleep following spatial experience. Science 271, 1870–1873 (1996).

34. Wilson, M. A. & McNaughton, B. L. Reactivation of hippocampal ensemble memories during sleep. Science 265, 676–679 (1994).

35. Yang, W. et al. Selection of experience for memory by hippocampal sharp wave ripples. Science 383, 1478–1483 (2024).

36. Davidson, T. J., Kloosterman, F. & Wilson, M. A. Hippocampal replay of extended experience. Neuron 63, 497–507 (2009).

37. Pfeiffer, B. E. & Foster, D. J. Hippocampal place-cell sequences depict future paths to remembered goals. Nature 497, 74–79 (2013).

38. Gillespie, A. K. et al. Hippocampal replay reflects specific past experiences rather than a plan for subsequent choice. Neuron 109, 3149–3163.e6 (2021).

39. Denovellis, E. L. et al. Hippocampal replay of experience at real-world speeds. Elife 10, (2021).

40. Mallory, C. S., Widloski, J. & Foster, D. J. The time course and organization of hippocampal replay. Science 387, 541–548 (2025).

41. Berners-Lee, A. et al. Hippocampal replays appear after a single experience and incorporate greater detail with more experience. Neuron 110, 1829–1842.e5 (2022).

42. Eichenbaum, H. Time cells in the hippocampus: a new dimension for mapping memories. Nat Rev Neurosci 15, 732–744 (2014).

43. Berndt, M., Trusel, M., Roberts, T. F., Pfeiffer, B. E. & Volk, L. J. Bidirectional synaptic changes in deep and superficial hippocampal neurons following in vivo activity. Neuron 111, 2984–2994.e4 (2023).

44. Norimoto, H. et al. Hippocampal ripples down-regulate synapses. Science 359, 1524–1527 (2018).

45. McClelland, J. L., McNaughton, B. L. & O’Reilly, R. C. Why there are complementary learning systems in the hippocampus and neocortex: insights from the successes and failures of connectionist models of learning and memory. Psychol Rev 102, 419–457 (1995).

46. Mankin, E. A. et al. Neuronal code for extended time in the hippocampus. Proc Natl Acad Sci U S A 109, 19462–19467 (2012).

47. Langston, R. F., Stevenson, C. H., Wilson, C. L., Saunders, I. & Wood, E. R. The role of hippocampal subregions in memory for stimulus associations. Behav Brain Res 215, 275–291 (2010).

48. Lee, J. S., Briguglio, J. J., Cohen, J. D., Romani, S. & Lee, A. K. The Statistical Structure of the Hippocampal Code for Space as a Function of Time, Context, and Value. Cell 183, 620–635.e22 (2020).

49. Wixted, J. T. et al. Sparse and distributed coding of episodic memory in neurons of the human hippocampus. Proc. Natl. Acad. Sci. U. S. A. 111, 9621–9626 (2014).

50. Olshausen, B. A. & Field, D. J. Emergence of simple-cell receptive field properties by learning a sparse code for natural images. Nature 381, 607–609 (1996).

51. El-Gaby, M. et al. An emergent neural coactivity code for dynamic memory. Nat. Neurosci. 24, 694–704 (2021).

52. Gava, G. P. et al. Organizing the coactivity structure of the hippocampus from robust to flexible memory. Science 385, 1120–1127 (2024).

53. Churchland, M. M. et al. Neural population dynamics during reaching. Nature 487, 51–56 (2012).

54. Churchland, M. M. & Shenoy, K. V. Preparatory activity and the expansive null-space. Nat Rev Neurosci 25, 213–236 (2024).

55. Libby, A. & Buschman, T. J. Rotational dynamics reduce interference between sensory and memory representations. Nat Neurosci 24, 715–726 (2021).

56. Huszár, R., Zhang, Y., Blockus, H. & Buzsáki, G. Preconfigured dynamics in the hippocampus are guided by embryonic birthdate and rate of neurogenesis. Nat Neurosci 25, 1201–1212 (2022).

57. Harvey, R. E., Robinson, H. L., Liu, C., Oliva, A. & Fernandez-Ruiz, A. Hippocampo-cortical circuits for selective memory encoding, routing, and replay. Neuron 111, 2076–2090.e9 (2023).

58. Soltesz, I. & Losonczy, A. CA1 pyramidal cell diversity enabling parallel information processing in the hippocampus. Nat Neurosci 21, 484–493 (2018).

59. Geiller, T., Fattahi, M., Choi, J.-S. & Royer, S. Place cells are more strongly tied to landmarks in deep than in superficial CA1. Nat Commun 8, 14531 (2017).

60. Lee, S.-H. et al. Parvalbumin-positive basket cells differentiate among hippocampal pyramidal cells. Neuron 82, 1129–1144 (2014).

61. Slomianka, L., Amrein, I., Knuesel, I., Sørensen, J. C. & Wolfer, D. P. Hippocampal pyramidal cells: the reemergence of cortical lamination. Brain Struct Funct 216, 301–317 (2011).

62. Valero, M. et al. Determinants of different deep and superficial CA1 pyramidal cell dynamics during sharp-wave ripples. Nat Neurosci 18, 1281–1290 (2015).

63. Spruston, N. Pyramidal neurons: dendritic structure and synaptic integration. Nat Rev Neurosci 9, 206–221 (2008).

64. Sharif, F., Tayebi, B., Buzsáki, G., Royer, S. & Fernandez-Ruiz, A. Subcircuits of Deep and Superficial CA1 Place Cells Support Efficient Spatial Coding across Heterogeneous Environments. Neuron 109, 363–376.e6 (2021).

65. Bilkey, D. K. & Schwartzkroin, P. A. Variation in electrophysiology and morphology of hippocampal CA3 pyramidal cells. Brain Res 514, 77–83 (1990).

66. Fitch, J. M., Juraska, J. M. & Washington, L. W. The dendritic morphology of pyramidal neurons in the rat hippocampal CA3 area. I. Cell types. Brain Res 479, 105–114 (1989).

67. Hemond, P. et al. Distinct classes of pyramidal cells exhibit mutually exclusive firing patterns in hippocampal area CA3b. Hippocampus 18, 411–424 (2008).

68. Hunt, D. L., Linaro, D., Si, B., Romani, S. & Spruston, N. A novel pyramidal cell type promotes sharp-wave synchronization in the hippocampus. Nat Neurosci 21, 985–995 (2018).

69. Magó, Á., Kis, N., Lükő, B. & Makara, J. K. Distinct dendritic Ca spike forms produce opposing input-output transformations in rat CA3 pyramidal cells. Elife 10, (2021).

70. Masukawa, L. M., Benardo, L. S. & Prince, D. A. Variations in electrophysiological properties of hippocampal neurons in different subfields. Brain Res 242, 341–344 (1982).

71. Raus Balind, S., et al. Diverse synaptic and dendritic mechanisms of complex spike burst generation in hippocampal CA3 pyramidal cells. Nat Commun 10, 1859 (2019).

72. Witter, M. P. Intrinsic and extrinsic wiring of CA3: indications for connectional heterogeneity. Learn Mem 14, 705–713 (2007).

73. Watson, J. F. et al. Human hippocampal CA3 uses specific functional connectivity rules for efficient associative memory. Cell 188, 501–514.e18 (2025).

74. Peng, Y., Barreda Tomás, F. J., Klisch, C., Vida, I. & Geiger, J. R. P. Layer-Specific Organization of Local Excitatory and Inhibitory Synaptic Connectivity in the Rat Presubiculum. Cereb Cortex 27, 2435–2452 (2017).

75. Peng, Y. et al. Directed and acyclic synaptic connectivity in the human layer 2-3 cortical microcircuit. Science 384, 338–343 (2024).

76. Douglas, R. J. & Martin, K. A. C. Neuronal circuits of the neocortex. Annu Rev Neurosci 27, 419–451 (2004).

77. Goodale, M. A. & Milner, A. D. Separate visual pathways for perception and action. Trends Neurosci 15, 20–25 (1992).

78. Mishkin, M., Ungerleider, L. G. & Macko, K. A. Object vision and spatial vision: two cortical pathways. Trends Neurosci. 6, 414–417 (1983).

79. Saur, D. et al. Ventral and dorsal pathways for language. Proc Natl Acad Sci U S A 105, 18035–18040 (2008).

80. Weiller, C., Reisert, M., Glauche, V., Musso, M. & Rijntjes, M. The dual-loop model for combining external and internal worlds in our brain. Neuroimage 263, 119583 (2022).

81. Knierim, J. J., Neunuebel, J. P. & Deshmukh, S. S. Functional correlates of the lateral and medial entorhinal cortex: objects, path integration and local-global reference frames. Philos Trans R Soc Lond B Biol Sci 369, 20130369 (2014).

82. Hargreaves, E. L., Rao, G., Lee, I. & Knierim, J. J. Major dissociation between medial and lateral entorhinal input to dorsal hippocampus. Science 308, 1792–1794 (2005).

83. Fernández-Ruiz, A. et al. Gamma rhythm communication between entorhinal cortex and dentate gyrus neuronal assemblies. Science 372, (2021).

84. Leutgeb, J. K., Leutgeb, S., Moser, M.-B. & Moser, E. I. Pattern separation in the dentate gyrus and CA3 of the hippocampus. Science 315, 961–966 (2007).

85. Rolls, E. T. The mechanisms for pattern completion and pattern separation in the hippocampus. Front Syst Neurosci 7, 74 (2013).

86. Gava, G. P. et al. Integrating new memories into the hippocampal network activity space. Nat Neurosci 24, 326–330 (2021).

87. Buzsáki, G. & Llinás, R. Space and time in the brain. Science 358, 482–485 (2017).

88. Vöröslakos, M., Petersen, P. C., Vöröslakos, B. & Buzsáki, G. Metal microdrive and head cap system for silicon probe recovery in freely moving rodent. Elife 10, (2021).

89. Angotzi, G. N. et al. Multi-Shank 1024 Channels Active SiNAPS Probe for Large Multi-Regional Topographical Electrophysiological Mapping of Neural Dynamics. Adv Sci (Weinh) 12, e2416239 (2025).

90. Osborne, J. E. & Dudman, J. T. RIVETS: a mechanical system for in vivo and in vitro electrophysiology and imaging. PLoS One 9, e89007 (2014).

91. Swanson, R. A. et al. Topography of putative bi-directional interaction between hippocampal sharp-wave ripples and neocortical slow oscillations. Neuron 113, 754–768.e9 (2025).

92. Vöröslakos, M. et al. 3D-printed Recoverable Microdrive and Base Plate System for Rodent Electrophysiology. Bio Protoc 11, e4137 (2021).

93. Siegle, J. H. et al. Open Ephys: an open-source, plugin-based platform for multichannel electrophysiology. J Neural Eng 14, 045003 (2017).

94. Mathis, A. et al. DeepLabCut: markerless pose estimation of user-defined body parts with deep learning. Nat Neurosci 21, 1281–1289 (2018).

95. Pachitariu, M., Sridhar, S., Pennington, J. & Stringer, C. Spike sorting with Kilosort4. Nat Methods 21, 914–921 (2024).

96. Petersen, P. C., Siegle, J. H., Steinmetz, N. A., Mahallati, S. & Buzsáki, G. CellExplorer: A framework for visualizing and characterizing single neurons. Neuron 109, 3594–3608.e2 (2021).

97. Watson, B. O., Levenstein, D., Greene, J. P., Gelinas, J. N. & Buzsáki, G. Network Homeostasis and State Dynamics of Neocortical Sleep. Neuron 90, 839–852 (2016).

98. Gonzalez, J., Torterolo, P. & Tort, A. B. L. Mechanisms and functions of respiration-driven gamma oscillations in the primary olfactory cortex. Elife 12, (2023).

99. Csicsvari, J., Jamieson, B., Wise, K. D. & Buzsáki, G. Mechanisms of gamma oscillations in the hippocampus of the behaving rat. Neuron 37, 311–322 (2003).

100. Semedo, J. D. et al. Feedforward and feedback interactions between visual cortical areas use different population activity patterns. Nat Commun 13, 1099 (2022).

101. Wainwright, M. J. High-Dimensional Statistics: A Non-Asymptotic Viewpoint. (Cambridge University Press, 2019).

102. González, J. et al. Low frequency oscillations drive EEG’s complexity changes during wakefulness and sleep. Neuroscience 494, 1–11 (2022).

103. Sebastian, E. R., Esparza, J. & M de la Prida, L. Quantifying the distribution of feature values over data represented in arbitrary dimensional spaces. PLoS Comput Biol 20, e1011768 (2024).

104. English, D. F. et al. Pyramidal Cell-Interneuron Circuit Architecture and Dynamics in Hippocampal Networks. Neuron 96, 505–520.e7 (2017).

